# NRF2-dependent regulation of the prostacyclin receptor PTGIR drives CD8 T cell exhaustion

**DOI:** 10.1101/2024.06.23.600279

**Authors:** Michael S. Dahabieh, Lisa M. DeCamp, Brandon M. Oswald, Susan M. Kitchen-Goosen, Zhen Fu, Matthew Vos, Shelby E. Compton, Joseph Longo, Kelsey S. Williams, Abigail E. Ellis, Amy Johnson, Ibukunoluwa Sodiya, Michael Vincent, Hyoungjoo Lee, Ryan D. Sheldon, Connie M. Krawczyk, Chen Yao, Tuoqi Wu, Russell G. Jones

## Abstract

The progressive decline of CD8 T cell effector function—also known as terminal exhaustion—is a major contributor to immune evasion in cancer. Yet, the molecular mechanisms that drive CD8 T cell dysfunction remain poorly understood. Here, we report that the Kelch-like ECH-associated protein 1 (KEAP1)-Nuclear factor erythroid 2-related factor 2 (NRF2) signaling axis, which mediates cellular adaptations to oxidative stress, directly regulates CD8 T cell exhaustion. Transcriptional profiling of dysfunctional CD8 T cells from chronic infection and cancer reveals enrichment of NRF2 activity in terminally exhausted (Tex^term^) CD8 T cells. Increasing NRF2 activity in CD8 T cells (via conditional deletion of KEAP1) promotes increased glutathione production and antioxidant defense yet accelerates the development of terminally exhausted (PD-1^+^TIM-3^+^) CD8 T cells in response to chronic infection or tumor challenge. Mechanistically, we identify PTGIR, a receptor for the circulating eicosanoid prostacyclin, as an NRF2-regulated protein that promotes CD8 T cell dysfunction. Silencing PTGIR expression restores the anti-tumor function of KEAP1-deficient T cells. Moreover, lowering PTGIR expression in CD8 T cells both reduces terminal exhaustion and enhances T cell effector responses (i.e. IFN-γ and granzyme production) to chronic infection and cancer. Together, these results establish the KEAP1-NRF2 axis as a metabolic sensor linking oxidative stress to CD8 T cell dysfunction and identify the prostacyclin receptor PTGIR as an NRF2-regulated immune checkpoint that regulates CD8 T cell fate decisions between effector and exhausted states.

**One Sentence Summary:** The KEAP1-NRF2 pathway is hyperactivated in terminally exhausted CD8 T cells and drives T cell dysfunction via transcriptional regulation of the prostacyclin receptor, *Ptgir*.

## INTRODUCTION

Immune evasion is a key hallmark of cancer. A common feature of many cancers is the progressive decline of anti-tumor CD8 T cell function that is critical for controlling tumor growth. CD8 T cell dysfunction—also known as T cell exhaustion—that is observed in cancer and chronic infections is driven by persistent exposure to antigens and suppressive factors in the tissue microenvironment (*1*). Features of exhausted T (Tex) cells include reduced proliferation, dampened production of effector molecules (i.e., IFN-γ, TNF-α, granzymes), and increased expression of inhibitory receptors that restrict T cell function (i.e., PD-1, CTLA-4, TIM-3) (*2*). Immune checkpoint inhibitors (ICIs) combat immune evasion by disrupting the activity of inhibitory receptors, thereby stimulating CD8 T effector (Teff) cell responses against tumor cells (*3*, *4*). While ICIs currently form the backbone of many cancer treatment regimens, clinical responses occur in only a subset of patients (*5*). Thus, a deeper understanding of the signaling axes that promote T cell dysfunction—which may extend beyond conventional protein-protein immune checkpoint interactions (i.e., PD-1/PD-L1)—is critical for identifying new ways to circumvent or reverse immune evasion in cancer.

While altered metabolism is a well-established hallmark of cancer (*6*), changes in cellular metabolism are also a core feature of dysfunctional T cells (*7*, *8*). Conventional CD8 Teff cells are powered by both glycolysis and oxidative phosphorylation (OXPHOS) (*9–11*), and use diverse fuels to drive anabolic metabolism and effector function (*12–14*). In contrast, CD8 Tex cells display impaired mitochondrial function and reduced OXPHOS (*15*, *16*). Oxidative stress, which can be triggered by mitochondrial dysfunction or metabolic conditions in the tumor microenvironment such as hypoxia, promotes the terminal exhaustion of CD8 T cells (*17*, *18*). Reducing levels of reactive oxygen species (ROS) produced during exposure to chronic antigen or hypoxia can prevent CD8 T cell terminal exhaustion (*17*, *18*). The antioxidant glutathione is essential for T cell metabolism and proliferation (*19*), establishing that management of cellular ROS levels is critical for productive CD8 T cell effector responses. While previous work has established mitochondrial dysfunction and oxidative stress as major factors limiting anti-tumor immunity, how T cells sense and manage oxidative stress to limit T cell dysfunction, particularly within the tumor microenvironment, remains poorly understood.

Here we identify the Nuclear factor erythroid 2-related factor 2 (NRF2) pathway, which is involved in sensing and responding to oxidative stress, as a key regulator of CD8 T cell terminal exhaustion. The Kelch-like ECH-associated protein 1 (KEAP1)-NRF2 signaling axis orchestrates the transcription of cytoprotective genes upon exposure to ROS or xenobiotics (*20*) and its activity is increased in terminally exhausted (Tex^term^) CD8 T cells. Leveraging a mouse model for conditional *Keap1* deletion in T cells (*CD4*^Cre^ *Keap1*^fl/fl^, referred to as *Keap1^−/−^*), we studied the impact of elevated NRF2 activity on CD8 T cell function. While increasing NRF2 activity enhanced glutathione production and reduced ROS levels in CD8 T cells, it paradoxically promoted CD8 T cell dysfunction, leading to accelerated terminal exhaustion in response to chronic infection or ectopic tumors. Mechanistically, we identified PTGIR, a G protein-coupled receptor (GPCR) for the eicosanoid lipid prostacyclin (*21*, *22*), as an NRF2-regulated protein that promotes CD8 T cell exhaustion. Ablating PTGIR expression restored the anti-tumor function of *Keap1^−/−^* T cells and was sufficient to enhance CD8 T cell effector function in response to chronic infection or cancer. These data establish the KEAP1-NRF2 axis as a metabolic sensor linking oxidative stress to CD8 T cell effector function and highlight the prostacyclin receptor PTGIR as an NRF2-regulated immune checkpoint that drives CD8 T cell terminal exhaustion.

## RESULTS

### NRF2 signaling promotes the terminal exhaustion of CD8 T cells

To investigate transcriptional changes associated with CD8 T cell exhaustion, we conducted a meta-analysis of RNA sequencing (RNA-seq) data from murine CD8 T cells isolated from *Listeria monocytogenes*- or lymphocytic choriomeningitis virus (LCMV)-infected mice or autochthonous liver cancer from published studies (*23–25*). Using relative gene expression levels of *bona fide* markers of naïve (Tn), effector (Teff), and exhausted (Tex) CD8 T cells **(Fig. 1A)**, we used unsupervised gene set enrichment analysis (GSEA) to identify pathways enriched in Tex versus Teff cells **(Fig. 1B, Table S1)**. Analysis of oncogenic signature gene sets (MSigDB C6 collection) revealed the NRF2 pathway as the most highly enriched dataset in Tex versus Teff cells (**Fig. 1B**). Mechanistically, under low or homeostatic ROS levels, KEAP1 dimers bind NRF2, promoting polyubiquitination of NRF2, which leads to its proteasomal degradation (**Fig. 1C**). Conversely, under high ROS levels, thiol residues on KEAP1 undergo reversible covalent modifications, resulting in conformational changes that ultimately liberate NRF2, leading to translocation of NRF2 to the nucleus and subsequent NRF2-dependent transcription of cytoprotective genes (*26*) **(Fig. 1C)**. NRF2 neutralizes ROS in part by promoting the transcription of enzymes involved in glutathione synthesis (i.e., *Gclc* (glutamate-cysteine ligase catalytic subunit) and *Gclm* (glutamate-cysteine ligase modifier subunit)) (*26*).

**Fig. 1.**
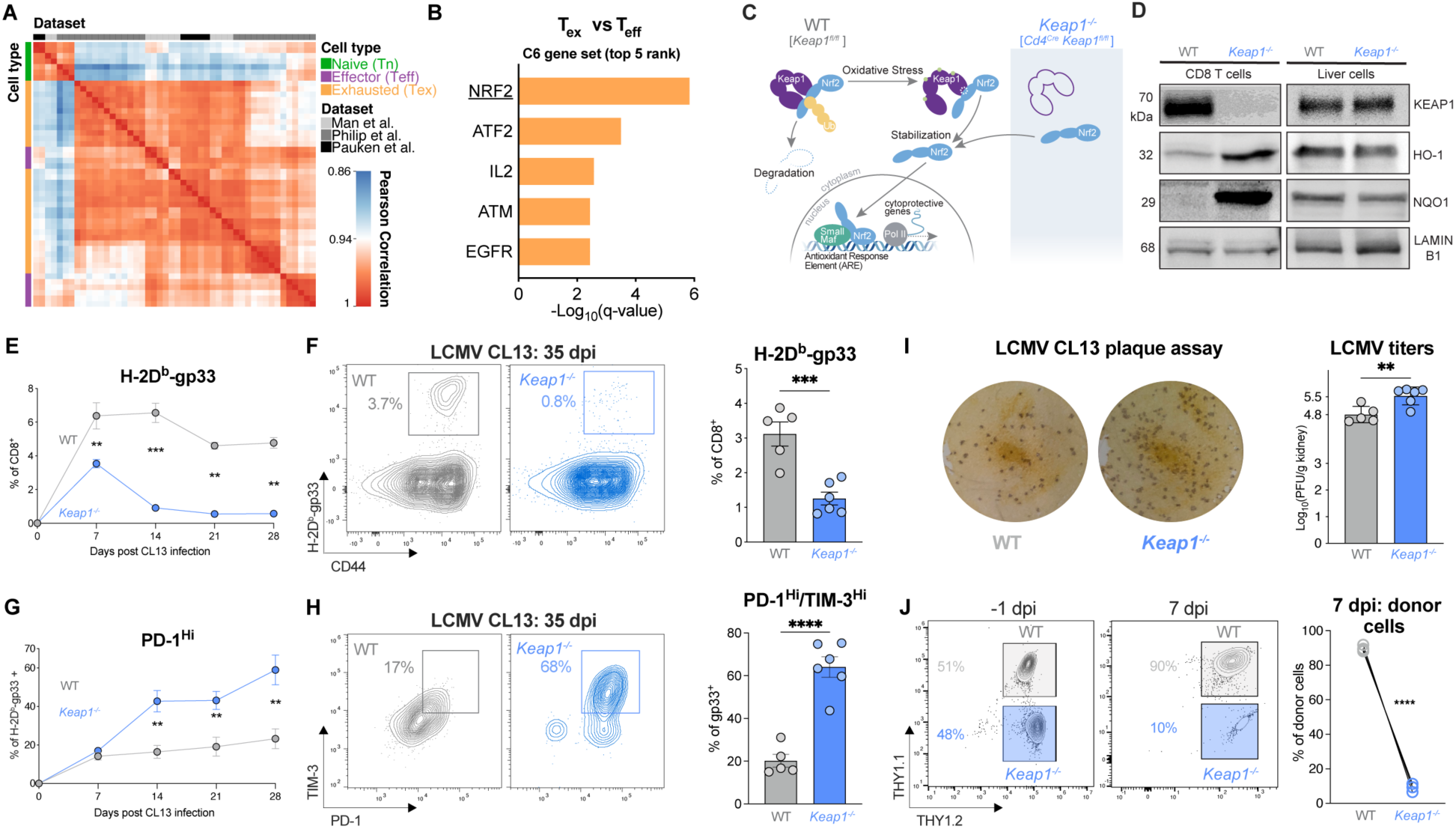
NRF2 signaling promotes CD8 T cell terminal exhaustion. **(A)** Pearson correlation plot of gene expression profiles for naïve (Tn), effector (Teff), and exhausted (Tex) CD8 T cells isolated from mice infected with *Listeria monocytogenes* or LCMV or mouse liver tumors. Data was obtained from published datasets (*23–25*). **(B)** Gene Set Enrichment Analysis (GSEA) of the top 5 oncogenic signature gene sets (MSigDB C6 gene set) enriched in Tex versus Teff cell clusters from (A). **(C)** Schematic of KEAP1-mediated regulation of NRF2 transcriptional activity. Graphic depicts KEAP1 binding and polyubiquitination of NRF2, leading to proteasomal degradation under homeostatic ROS levels. Oxidative stress promotes modification of thiol residues on KEAP1, leading to translocation of NRF2 to the nucleus and activation of NRF2-dependent transcription. Deletion of *Keap1* in T cells (*CD4^Cre^ Keap1^fl/fl^*) promotes NRF2 translocation to the nucleus, independent of oxidative stress. **(D)** Immunoblot of KEAP1, NRF2 transcriptional targets (HO-1 and NQO1), and LAMIN B1 (loading control) in activated CD8 T cells and liver cells from WT and T cell *Keap1*-deficient (*Keap1^−/−^*) mice. **(E-F)** Percentage of LCMV-specific CD8 T cells in WT and *Keap1*^−/−^ mice following LCMV clone 13 (CL13) infection. Shown is the percentage of H-2D^b^-gp33 tetramer-binding CD8 T cells in (E) the blood over time or (F) the spleen of mice at 35 days post infection (dpi) as determined by flow cytometry. Bar graph shows the percentage of gp33-specific CD8 T cells in the spleen of infected mice at 35 dpi (mean±SEM, n=5-6). **(G-H)** Inhibitory receptor expression on H-2D^b^-gp33 tetramer-positive CD8 T cells following LCMV CL13 infection of WT and *Keap1*^−/−^ mice. Shown are (G) the percentage of PD-1^Hi^ gp33-specific CD8 T cells over the course of LCMV CL13 infection and (H) the percentage of Tex^term^ (PD-1^HI^TIM-3^HI^) gp33-specific CD8 T cells in WT versus *Keap1*^−/−^ mice at 35 dpi. Bar graph shows the percentage of PD-1^HI^TIM-3^HI^ gp33-specific CD8 T cells in the spleen at 35 dpi (mean±SEM, n=5-6). **(I)** LCMV CL13 plaque assay from kidney extracts of WT and *Keap1*^−/−^ mice at 35 dpi. Shown are representative plaque images and bar graph quantifying the number of plaque forming units (PFU) per gram of tissue (mean±SEM, n=5-6). **(J)** Expansion of WT and *Keap1*^−/−^ P14 CD8 T cells following co-transfer into naïve hosts and infection with LCMV CL13. Shown are flow cytometry plots depicting the percentage of WT and *Keap1*^−/−^ P14 T cells at the time of transfer (−1 dpi) and on 7 dpi with LCMV CL13, with quantification per genotype shown by the line graph (mean±SEM, n=4. \*\**P*<0.01, \*\*\**P*<0.001, \*\*\*\**P*<0.0001.

To investigate the role of NRF2 signaling in CD8 T cell responses, we generated a mouse model with conditional deletion of *Keap1* in mature T cells using *Cd4^Cre^* transgenic mice (**Fig. 1C**). RT-PCR analyses confirmed knockout of *Keap1* (exon 2), while NRF2 transcriptional targets NAD(P)H quinone dehydrogenase 1 (*Nqo1*), Heme oxygenase-1 (*Hmox1*), and *Gclc* were transcriptionally elevated in CD8 T cells from T cell KEAP1-deficient (*Keap1*^−/−^) versus control (WT) mice **(Fig. S1A, Table S2)**. Similarly, immunoblot of KEAP1 confirmed its deletion in *Keap1^−/−^* T cells, while the protein products of NRF2 transcriptional targets *Hmox1* (HO-1) and *Nqo1* (NQO1) were upregulated in *Keap1^−/−^* T cells **(Fig. 1D)**. To validate that *Keap1* deletion was specific to T cells, we confirmed that hepatocytes from *Keap1*^fl/fl^ (WT) and *Cd4^Cre^ Keap1^fl/fl^* (*Keap1*^−/−^) mice displayed normal expression of NRF2 target genes at both the transcriptional **(Fig. S1B)** and protein (**Fig. 1D**) level. Deletion of *Keap1* in the T cell lineage led to a modest reduction in the percentage of CD3^+^ T cells in *Keap1*^−/−^ versus WT mice **(Fig. S2A)**, but no significant difference in the percentage of CD8^+^ T cells between genotypes **(Fig. S2B)**. T cell *Keap1*-deficient mice displayed decreased percentages and numbers of effector and central memory CD8^+^ T cells **(Fig. S2C)**, reduced effector CD4^+^ T cells **(Figs. S2D-E)**, elevated percentages of B cells and natural killer cells, and decreased populations of NKT cells **(Fig. S2F)**. Innate cells including neutrophils **(Fig. S2G)**, and conventional type I dendritic cells (cDC1) were also reduced in *Keap1^−/−^* compared to WT mice **(Fig. S2H)**.

Our data confirm that loss of KEAP1 promotes hyperactivation of NRF2 in CD8 T cells (**Fig. 1D**), providing a suitable model to examine the role of NRF2 activation in the context of CD8 T cell exhaustion. To test this, we challenged control (WT) and *Keap1* T cell-deficient (*Keap1*^−/−^) mice with the clone 13 (CL13) strain of LCMV, which induces chronic infection and the emergence of PD-1^+^ CD8 Tex cells in vivo (*2*). We first assessed the percentage of antigen specific (H-2D^b^-gp33^+^-binding) CD8 T cells in the blood of WT and *Keap1*^−/−^ mice over the course of CL13 infection. As early as 7 days post infection (dpi), we observed a significant reduction in the percentage of circulating H-2D^b^-gp33-specific CD8 T cells in *Keap1*^−/−^ mice compared to controls (~3% vs 6%), which was sustained over the course of infection **(Fig. 1E)**. Similarly, we observed a significant reduction in the percentage of gp33-specific CD8 T cells in the spleen of *Keap1*^−/−^ (~1%) versus WT mice (~3%) at 35 dpi **(Fig. 1F).** Strikingly, we observed a marked increase in PD-1 expression on antigen-specific *Keap1*^−/−^ T cells compared to controls as early as 14 days post LCMV CL13 infection, which was maintained throughout the course of infection **(Fig. 1G)**. At 35 dpi, the majority of gp33-specific CD8 T cells from *Keap1*^−/−^ mice displayed elevated expression of both PD-1 and T cell immunoglobulin and mucin domain-containing protein 3 (TIM-3)—surface markers of terminal exhaustion—which was elevated over three-fold compared to controls (~65% for *Keap1*^−/−^ versus ~20% for WT, **Fig. 1H**). Thus, hyperactivation of NRF2 (driven by loss of KEAP1) leads to a defect in the expansion and persistence of antigen-specific CD8 T cells in response to chronic infection and promotes CD8 T cell differentiation towards a terminally exhausted (Tex^term^) state.

We next assessed the functional consequence of NRF2 hyperactivation during chronic infection by quantifying viral loads in the kidneys of CL13-infected mice. We observed a significant increase in LCMV plaque forming units (PFU) in *Keap1*^−/−^ (Log_10_ 5.5 PFU) compared to WT (Log_10_ 4.8 PFU) mice **(Fig. 1I)**, which correlated with increased numbers of terminally exhausted (PD-1^+^TIM-3^+^) CD8 T cells in *Keap1*^−/−^ mice **(Figs. 1G-H)**. Finally, to establish whether the effects of KEAP1 loss in CD8 T cells were cell-intrinsic, we co-transferred control (WT) and *Keap1^−/−^* P14 CD8 T cells into congenic hosts, followed by challenge with LCMV CL13. In this competitive model, we observed a marked reduction in the expansion of *Keap1^−/−^*P14 T cells relative to control P14 T cells in vivo, indicating that the poor expansion of CD8 T cells in T cell *Keap1*-deficient mice was intrinsic to CD8 T cells **(Fig. 1J)**. Collectively, our data indicate that loss of KEAP1 promotes CD8 T cell exhaustion, as shown by reduced T cell expansion, increased features of terminal exhaustion, and increased viral burden during chronic viral infection.

### NRF2 activation is linked to CD8 T cell dysfunction in cancer and poor tumor control

We next examined the impact of NRF2 hyperactivation on CD8 tumor infiltrating lymphocyte (TIL) function and tumor clearance. We first examined NRF2 pathway activity in CD8 TIL populations sorted from B16 melanoma tumors grown in C57BL/6 mice and analyzed by single cell RNA sequencing (scRNA-seq). A Uniform Manifold Approximation and Projection (UMAP; (*27*)) of cell-specific transcripts identified five major CD8^+^ T cell populations in B16 tumors based on previously reported markers (*25*, *28*): proliferating (*Cdk1*^hi^), progenitor exhausted (Tex^prog^), PD-1^+^LAG3^+^ cells with expression of killer cell lectin-type receptors (Tex^KLR^) or interferon-stimulated genes (Tex^ISG^), and PD-1^+^TIM-3^+^ terminally exhausted (Tex^term^, *Tox*^hi^) T cells (**Fig. 2A, Fig. S3A**). Next we conducted GSEA for NRF2 signatures (*29*, *30*). This analysis revealed that *Tox*^hi^ Tex^term^ TIL expressed the highest NRF2 signature and pathway activity score compared to all other clusters **(Fig 2B-C, Fig. S3B)**, indicating that elevated NRF2 activity is associated with CD8 T cell terminal exhaustion.

**Fig. 2.**
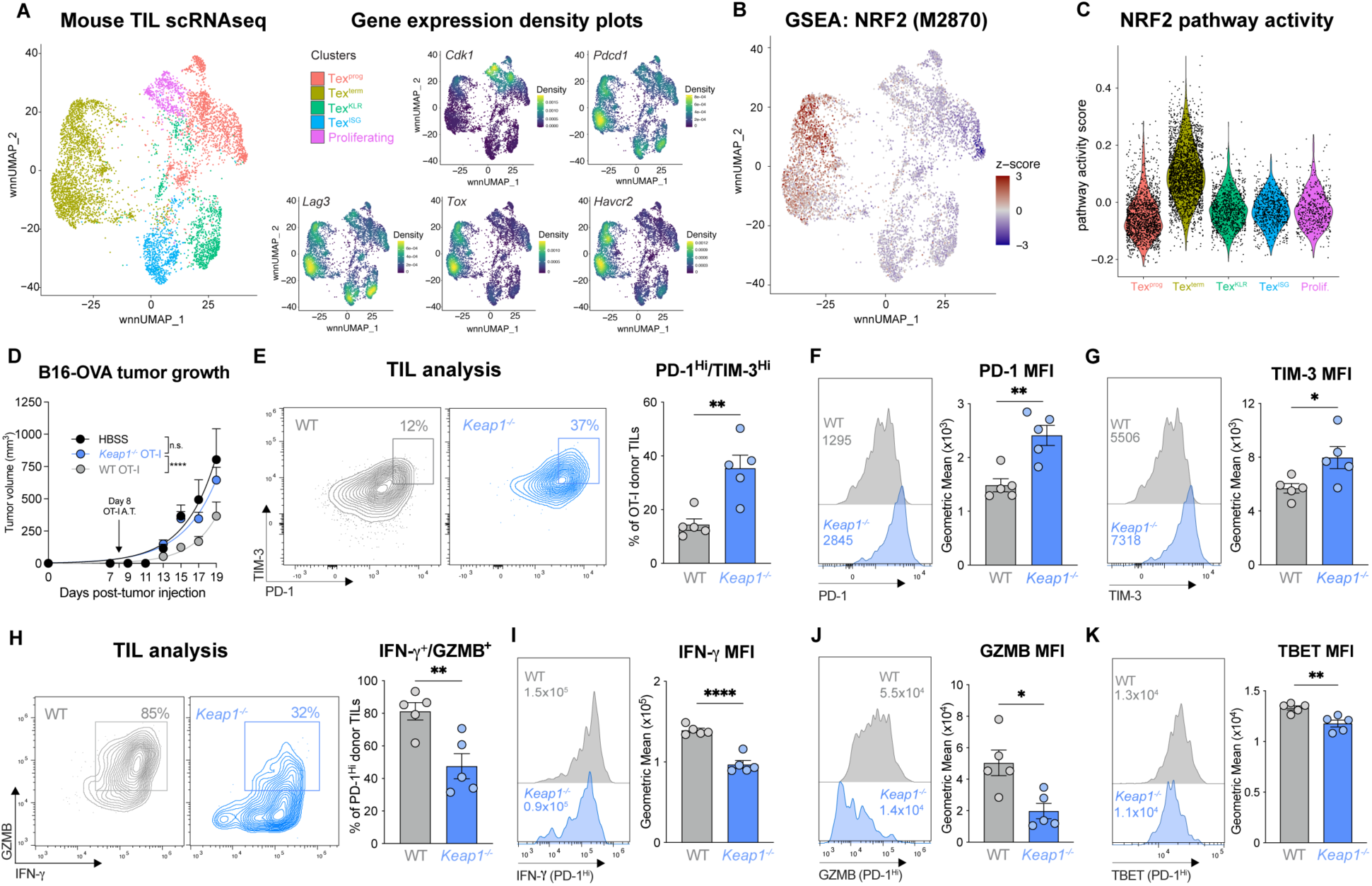
NRF2 activation is linked to CD8 T cell dysfunction in cancer and poor tumor control. **(A)** Weighted nearest neighbor Uniform Manifold Approximation and Projection (wnn_UMAP) of cell-specific transcripts for the 5 major activated (CD44^+^) CD8^+^ T cell clusters in B16 tumors (14 days post tumor inoculation (dpti)): progenitor exhausted (Tex^prog^), terminally exhausted (Tex^term^), exhausted killer cell lectin-like receptor-expressing (Tex^KLR^), exhausted IFN-I stimulated gene expressing (Tex^ISG^), and proliferating (prolif.). Gene expression density plots for *Cdk1, Pdcd1, Lag3, Tox,* and *Havcr2* are shown. **(B)** GSEA embedding of NRF2 transcriptional target signature (M2870) in the CD8 TIL UMAP from (A). **(C)** Violin plot of NRF2 (M2870) pathway activity score for each of CD8 T cell clusters defined in (A). **(D)** B16-OVA melanoma tumor growth in mice following adoptive transfer of WT or *Keap1^−/−^* OT-I CD8 T cells at 8 dpti (mean±SEM, n=11. **(E)** PD-1 versus TIM-3 expression on WT and *Keap1^−/−^* OT-I CD8 TIL isolated from B16 tumors at 15 dpti (mean±SEM, n=5. **(F-G)** Histogram of (F) PD-1 and (G) TIM-3 expression on WT or *Keap1^−/−^* OT-I CD8 TIL isolated from B16 tumors as in (E), with geometric mean fluorescence intensity (gMFI) shown in inset. Bar graphs quantify the gMFI of PD-1 and TIM-3 expression (mean±SEM, n=5. **(H)** Representative flow cytometry plot of IFN-γ versus Granzyme B (GZMB) expression for PD-1^HI^ WT and *Keap1^−/−^* TIL isolated as in (E). **(I-K)** Representative histograms and bar graph of gMFI for (I) IFN-γ, (J) GZMB, and (K) TBET expression by PD-1^HI^ WT or *Keap1^−/−^* OT-I CD8 TIL isolated from B16 tumors as in (E). \**P*<0.05, \*\**P*<0.01, \*\*\**P*<0.001, \*\*\*\**P*<0.0001.

Given the pronounced NRF2 signature displayed by Tex^term^ cells in tumors (**Fig. 2B-C**) and the increased features of exhaustion shown by *Keap1^−/−^* CD8 T cells during chronic LCMV infection **(Fig. 1)**, we hypothesized that NRF2 hyperactivation may promote CD8 T cell exhaustion in tumors. To test this, we first inoculated *Cd4^Cre^ Keap1^fl/fl^*(*Keap1*^−/−^) mice and *Keap1^fl/fl^* controls (WT) with MC38 colorectal cancer cells subcutaneously in their rear flank and tracked tumor growth over time. We observed accelerated tumor growth in T cell-specific KEAP1-deficient mice compared to controls as determined by faster time to tumor onset and experimental endpoint (tumor size >1500 mm^3^) (**Fig. S3C**). Notably, by day 40, all *Keap1*^−/−^ animals had reached experimental endpoint, while ~30% of control mice did not develop tumors **(Fig. S3C).** We next assessed the impact of KEAP1 deficiency on T cell-intrinsic control of tumor growth via adoptive transfer of tumor-specific CD8 T cells using a second tumor model, ovalbumin (OVA)-expressing B16 melanoma cells. CD45.1^+^ C57BL/6 mice inoculated with B16-OVA tumors received either WT or *Keap1*^−/−^ OVA-specific OT-I CD8 T cells (CD45.2^+^) 8 days post tumor inoculation. As expected, B16-OVA tumor growth was significantly reduced in mice that received WT OT-I T cells compared to HBSS-injected vehicle control mice; however, mice that received *Keap1*^−/−^ OT-I T cells displayed similar rates of tumor growth as mice that received no OT-I T cells **(Fig. 2D, Fig. S3D)**.

We next assessed the impact of elevated NRF2 activity on the anti-tumor functions of CD8 T cells. TIL were isolated from a cohort of B16-OVA tumor-bearing mice that received either WT or *Keap1*^−/−^ OT-I T cells as in **Fig. 2D**. We observed a ~3-fold increase in the number of terminally exhausted (PD-1^hi^TIM-3^hi^) OT-I T cells in B16 tumors when KEAP1 was deleted **(Fig. 2E)**. *Keap1^−/−^* OT-I TIL from tumor-bearing mice also displayed higher PD-1 and TIM-3 protein expression compared to controls, as determined by a shift in the mean fluorescence intensity (MFI) of these markers **(Fig. 2F-G).** Functionally, we observed lower IFN-γ- and Granzyme B (GZMB)-expression in OT-I TIL from B16-OVA tumors when KEAP1 was deleted **(Fig. 2H)**. Focusing on the PD-1^+^ TIL subset, OT-I TIL lacking KEAP1 expression displayed lower levels of IFN-γ (**Fig. 2I**), GZMB (**Fig. 1J**), and TBET (**Fig. 2K**) proteins relative to control OT-I TIL, consistent with overall reduced function due to KEAP1 loss. Together, these data indicate that elevated NRF2 activity (due to KEAP1 loss) promotes CD8 T cell dysfunction in tumors, reducing T cell-intrinsic control of tumor growth in vivo.

### NRF2 promotes antioxidant defense but also drives CD8 T cell exhaustion

NRF2 binding is a hallmark interaction for KEAP1 (*31*); however, KEAP1 also interacts with other regulators of cellular metabolism, including the autophagy receptor p62 (*32*, *33*) and the glycolytic enzyme glyceraldehyde 3-phosphate dehydrogenase (GAPDH) (*34*). To establish whether NRF2 is the downstream target of KEAP1 loss leading to CD8 T cell dysfunction, we genetically ablated NRF2 (encoded by *Nfe2l2*) in primary KEAP1-deficient CD8 T cells using single guide (sg) RNAs targeting *Nfe2l2* **(Table S2)** (*35*). Proteomics analysis validated the depletion of NRF2 in *Keap1^−/−^* CD8 T cells that received *Nfe2l2*-targeting sgRNA (*sgNfe2l2*) but not scrambled control (*sgScr*) (**Fig. 3A, Fig. S4A**). NRF2 transcriptional activity was reduced in *Nfe2l2*-deleted *Keap1^−/−^* CD8 T cells as evidenced by reduced protein expression of GCLM—an enzyme essential for glutathione synthesis—which is transcriptionally regulated by NRF2 (*36–38*) **(Fig. 3A, Fig. S4A)**.

**Fig. 3.**
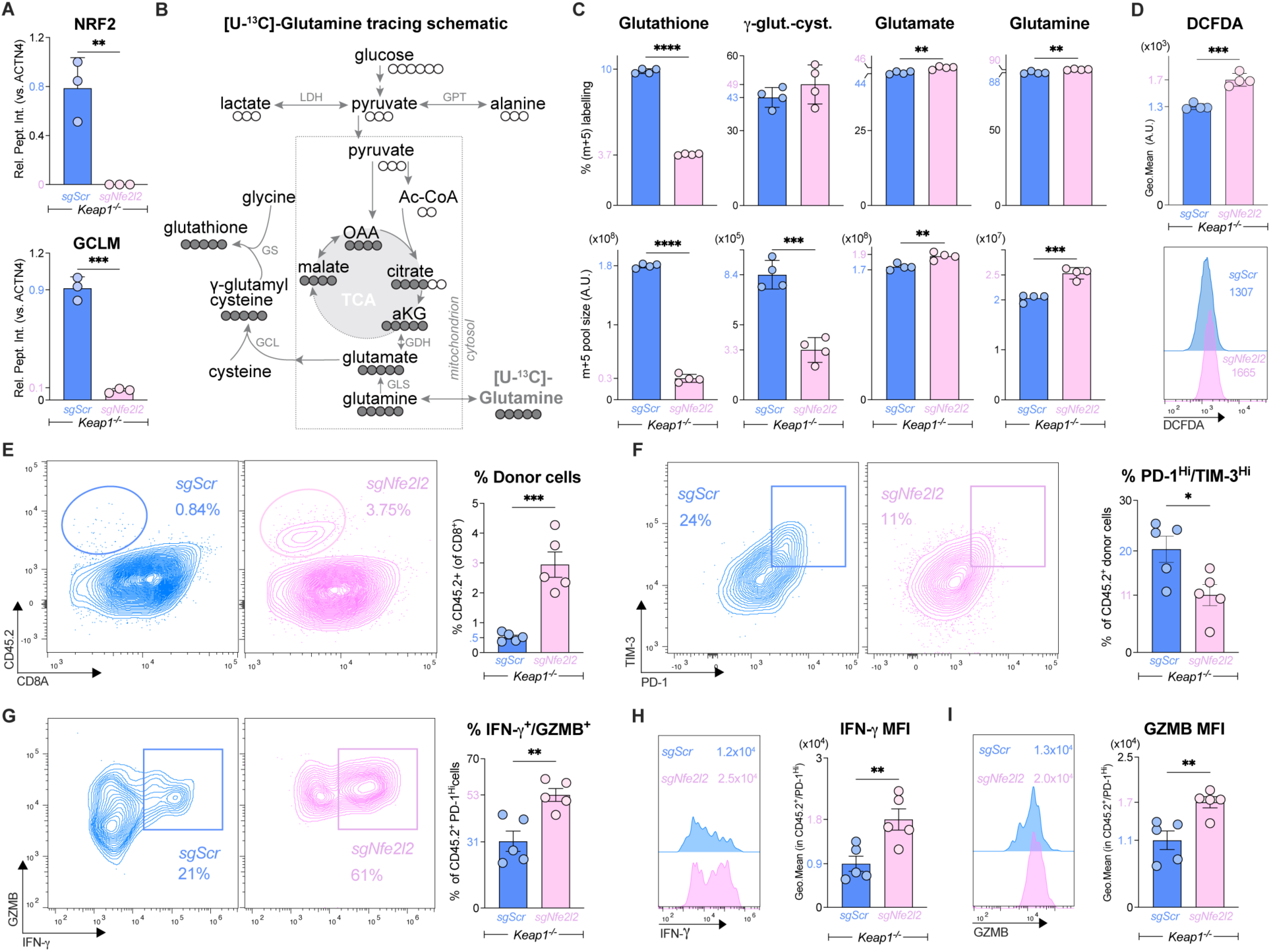
NRF2 promotes antioxidant defense but also drives CD8 T cell exhaustion. **(A)** Relative abundance of NRF2 and GLCM protein in CRISPR/Cas9 edited *Keap1^−/−^* CD8^+^ P14 T cells that received *Nfe2l2*-targeting (*sgNfe2l2*) or control (*sgScr*) sgRNAs as determined by proteomics (mean±SEM, n=3/sample). Protein levels are expressed relative to ACTN4 levels. **(B)** Schematic of glutathione synthesis from [U-^13^C]-glutamine. [U-^13^C]-glutamine-labeled carbon is shown in closed circles. Enzymes involved in glutathione synthesis from glutamine are shown: glutaminase (GLS), glutamate cysteine ligase (GCL), and glutathione synthase (GS). Also shown are Lactate Dehydrogenase (LDH) and Glutamic-Pyruvic Transaminase (GPT) and integration of glutamine into the TCA cycle via Glutamate Dehydrogenase (GDH). **(C)** Bar graphs depicting percentage of m+5 isotopologue labeling (top) and total m+5 labeled pool size (bottom) for glutathione, γ-glutamyl cysteine (γ-glut-cyst), glutamate, and glutamine in *sgNfe2l2* and *sgScr* edited *Keap1^−/−^*CD8 T cells (mean±SEM, n=4/sample). **(D)** Representative histogram and bar graphs quantifying DCFDA fluorescence in *sgNfe2l2* and *sgScr* edited *Keap1^−/−^*CD8 T cells (mean±SEM, n=4/sample). **(E)** Representative flow cytometry plots of CD8 versus CD45.2 expression on CD8^+^ gated splenocytes harvested from LCMV CL13-infected mice at 7 dpi that received *sgNfe2l2* and *sgScr* edited *Keap1^−/−^* P14 T cells at −1 dpi. CD45.2 marks adoptively transferred P14 T cells. Bar graphs represent the percentage of CD45.2^+^ P14 T cells of total CD8^+^ splenocytes (mean±SEM, n=5/sample). **(F)** Representative flow cytometry plots of PD-1 versus TIM-3 expression by donor CD45.2^+^ *sgNfe2l2* and *sgScr Keap1^−/−^* P14 T cells at 7 dpi from (E). Bar graph quantifies the percent PD-1^Hi^/TIM-3^Hi^ CD45.2^+^ *Keap1^−/−^* P14 T cells in the spleen (mean±SEM, n=5/sample). **(G-I)** Representative flow cytometry plot of IFN-γ and GZMB by PD-1^Hi^ *sgNfe2l2* and *sgScr Keap1^−/−^* P14 T cells at 7 dpi as in (E). Bar graph quantifies the polyfunctional (IFN-γ^+^GZMB^+^) CD45.2^+^ *Keap1^−/−^* P14 T cells in the spleen (mean±SEM, n=5/sample). **(H-I)** Representative histograms and bar graph of gMFI for (H) IFN-γ and (I) GZMB expression by PD-1^Hi^ donor cells from (E) at 7 dpi (mean±SEM, n=5/sample). \**P*<0.05, \*\**P*<0.01, \*\*\**P*<0.001, \*\*\*\**P*<0.0001.

We next assessed the impact of silencing NRF2 on the metabolism of *Keap1^−/−^* CD8 T cells. A major cytoprotective function of NRF2 is transcriptional regulation of antioxidant defense systems, including glutathione production (*39*, *40*). Glutathione is synthesized *de novo* from glutamate, cysteine, and glycine (**Fig. 3B**). NRF2 increases glutathione synthesis through transcriptional upregulation of components of the glutamate cysteine ligase (GCL) complex—a heterodimer composed of catalytic (GCLC) and modifier (GCLM) subunits—which mediates the condensation of glutamate with cysteine to form γ-glutamyl cysteine (**Fig. 3B**) (*41*). We used metabolomics to track the production of glutathione from uniformly labelled [U]-^13^C-glutamine in *Keap1^−/−^* T cells lacking NRF2 (*sgNfe2l2*) compared to control (*sgScr*) *Keap1^−/−^* T cells. Silencing NRF2 led to a slight increase in intracellular glutamine in *Keap1^−/−^* T cells (likely from increased uptake of [U]-^13^C-glutamine) (**Fig. S4B**) but had no impact on TCA cycle metabolism (**Fig. S4C**). The major metabolic impact of silencing NRF2 in *Keap1^−/−^* T cells was a ~3-fold reduction in glutathione levels, as determined by both the percentage and pool size (intensity) of m+5-labelled glutathione from [U]-^13^C-glutamine (**Fig. 3C, Fig. S4D**). This reduction in glutathione production was linked to reduced GCL activity, as GCLM protein levels were decreased ~90% (**Fig. 3A**) and m+5 γ-glutamyl cysteine production from [U]-^13^C-glutamine was reduced >60% in NRF2-deficient *Keap1^−/−^* T cells (**Fig. 3C, Fig. S4D**). Since glutathione serves as an antioxidant in its reduced form, we measured reactive oxygen species (ROS) levels in *Keap1^−/−^* T cells using dichlorodihydrofluorescein diacetate (DCFDA), a fluorogenic reporter for intracellular ROS (*42*). Consistent with reduced glutathione synthesis from [U]-^13^C-glutamine, NRF2-deficient *Keap1^−/−^* T cells displayed a 50% increase in DCFDA staining compared to controls **(Fig. 3D)**, indicative of increased levels of oxidative stress. Thus, NRF2 drives the metabolic program of glutathione synthesis (via GCLM) that regulates ROS levels in CD8 T cells.

We next assessed the impact of silencing NRF2 on the response of *Keap1^−/−^*T cells to chronic infection with LCMV CL13. Control (*sgScr)* and NRF2 knockout (*sgNfe2l2*) P14 *Keap1^−/−^* T cells (CD45.2^+^ background) were transferred into congenic (CD45.1^+^) hosts one day prior to infection with LCMV CL13. At 7 dpi, flow cytometry analysis of splenic lymphocytes revealed a robust ~6-fold increase in the percentage and number of NRF2 knockout donor P14 *Keap1^−/−^* T cells compared to *Keap1^−/−^* controls **(Fig. 3E, S5A)**. The rescue in *Keap1^−/−^* T cell expansion in vivo by silencing NRF2 was accompanied by both a reduction in inhibitory receptor expression and a restoration of T cell effector function. Mice that received NRF2-deficient P14 *Keap1^−/−^* T cells displayed a 50% reduction in the percentage of terminally exhausted (PD-1^+^TIM-3^+^) P14 T cells in the spleen **(Fig. 3F, S5B)**, while NRF2 knockout PD-1^hi^ T cells displayed a 2-fold increase in the percentage and 12-fold increase in the number of IFN-γ- and GZMB-producing cells in response to LCMV CL13 infection **(Fig. 3G, Figs. S5C-E)**. Silencing NRF2 in *Keap1^−/−^* P14 T cells led to a marked shift in both IFN-γ **(Fig. 3H)** and GZMB **(Fig. 3I)** protein production, even in cells expressing elevated levels of PD-1. While the percentage of TNF-α-producing P14 T cells was similar between control (*sgScr*) and NRF2-deleted (*sgNfe2l2*) *Keap1^−/−^* P14 T cells **(Fig. S5F)**, due to the restored expansion conferred by NRF2 deletion we observed an overall increase in the number of TNF-α-producing *Keap1^−/−^* P14 T cells in the spleen upon NRF2 silencing **(Fig. S5F,G).**

Finally, we examined the expression of transcription factors TOX, TCF1 and TBET, which orchestrate the transition from progenitor exhausted to terminally exhausted states (*43*), in *Keap1^−/−^* P14 T cells following LCMV CL13 infection. PD-1^hi^ NRF2 knockout *Keap1^−/−^* P14 T cells displayed a shift in TCF1 expression and no change in TBET expression compared to by control *Keap1^−/−^* P14 T cells **(Figs. S5H-K)**. This increase in TCF1 expression resulted in higher percentage and number of TOX^+^TCF1^+^ P14 T cells when NRF2 was deleted in *Keap1^−/−^* T cells **(Figs. S5H)**, consistent with a shift in the PD-1^hi^ population away from a TOX+ Tex^term^ state and towards a quiescent resident/proliferating circulating progenitor state (*43*). Collectively, these data establish that hyperactivation of the NRF2 pathway drives CD8 T cells towards a terminally exhausted state, despite providing metabolic adaptations to CD8 T cells (i.e., increasing glutathione production) that reduce ROS levels.

### NRF2 directly regulates expression of the Prostacyclin receptor (*Ptgir*)

To interrogate how NRF2 hyperactivation drives CD8 T cell exhaustion, we analyzed changes in mRNA levels in *Keap1^−/−^* versus WT P14 cells following LCMV infection. WT or *Keap1^−/−^* P14 T cells (CD45.2^+^) were adoptively transferred into congenic CD45.1^+^ hosts followed by infection with LCMV, and P14 T cells were sorted via flow cytometry at 7 dpi for RNA-seq analysis (**Fig. 4A**). Differential gene expression analysis verified that hallmark NRF2 targets including *Nqo1* and *Gclc* were upregulated in *Keap1^−/−^* P14 T cells compared to controls, while WT cells displayed high expression levels of cell-division associated genes and killer cell lectin-like receptor family genes: *Klra3, Klra4, Klrk1* **(Fig. 4B, Table S3)**. Pathway enrichment analysis identified NRF2 targets and oxidative stress response pathways as signatures of *Keap1^−/−^* P14 T cells, consistent with hyperactivation of NRF2 in these T cells **(Figs. S6A-D, Table S4)**. Notably, the most upregulated transcript in *Keap1^−/−^* P14 cells was *Ptgir* **(Fig. 4B)**, which encodes for the receptor that binds prostacyclin— a circulating eicosanoid family lipid that is widely characterized to bind PTGIR on vascular smooth muscle cells, resulting in vasodilation and platelet aggregation. (*44*). Prostacyclin is initially synthesized via Phospholipase A2-mediated liberation of arachidonic acid (AA) from cell membrane phospholipids. AA then serves as a substrate for cyclooxgenase-1 and -2 (COX-1 and COX-2) to synthesize prostaglandin H2 (PGH_2_), which is isomerized to prostacyclin (PGI_2_) via prostacyclin synthase (PTGIS) (*45*). Using qPCR, we validated that *Ptgir* mRNA expression was increased >1000-fold in *Keap1^−/−^* versus control P14 CD8^+^ T cells responding to LCMV infection in vivo **(Fig. 4C)**. The relative induction of *Ptgir* mRNA in *Keap1^−/−^* P14 T cells in response to LCMV infection was greater than the conventional NRF2 target *Nqo1*, which was increased ~50-fold over control T cells **(Fig. 4C)**.

**Fig. 4.**
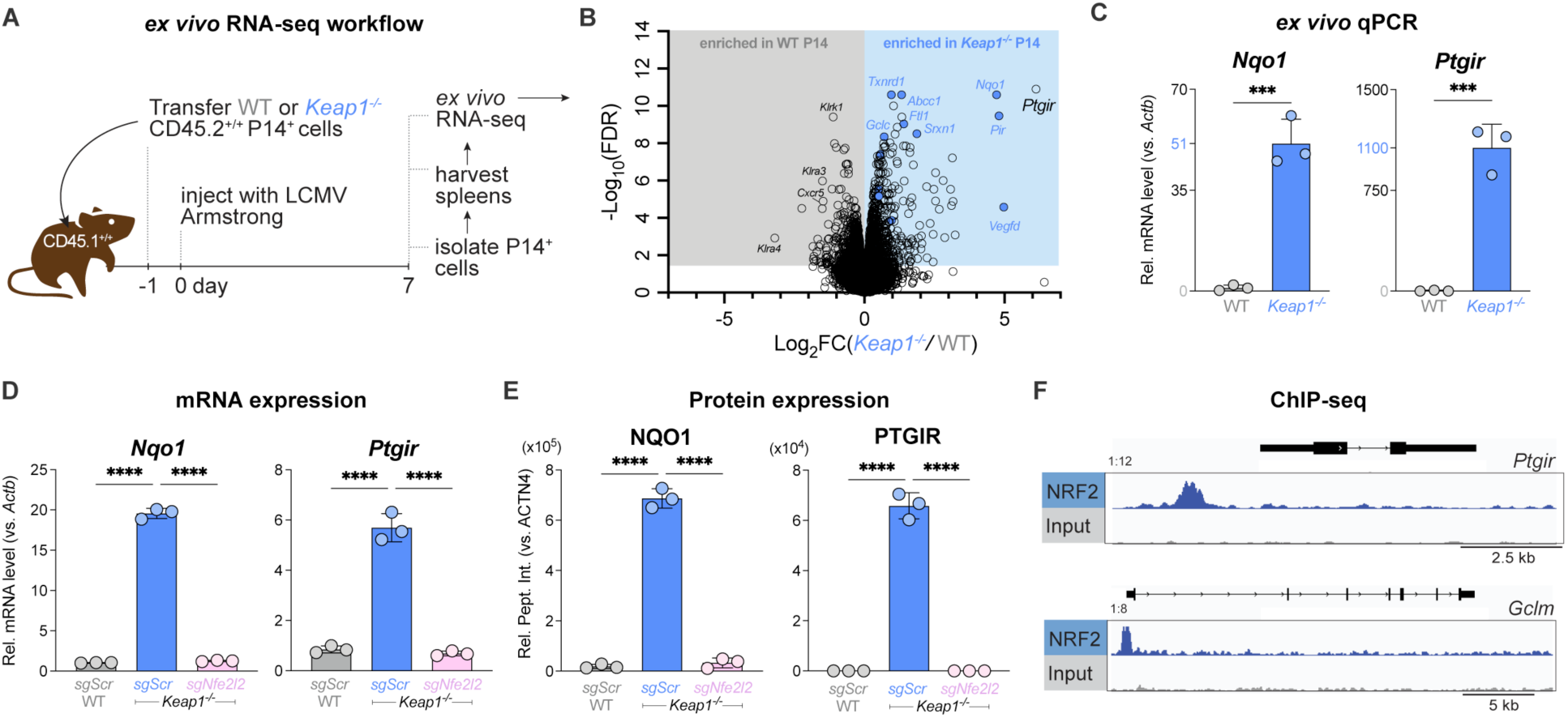
NRF2 directly regulates expression of the Prostacyclin receptor (*Ptgir*). **(A)** Schematic of RNA-seq workflow for P14 T cells responding to LCMV infection. WT or *Keap1^−/−^* P14 T cells (CD45.2^+^) were adoptively transferred into congenic CD45.1^+^ hosts followed by infection with LCMV (Armstrong strain). At 7 dpi, P14 T cells were sorted via CD45.2 positive selection for bulk RNA-seq analysis (3 biological replicates/genotype). **(B)** Volcano plot (horizontal axis: Log_2_ fold change (*Keap1^−/−^*/WT), vertical axis: −Log_10_ False Discovery Rate (FDR)) highlighting differentially expressed genes (DEGs) in WT and *Keap1^−/−^* P14 T cells at 7 dpi. Gray and blue shaded regions represent genes significantly enriched (-Log_10_ FDR > 1.30, corresponding to FDR <0.05) in WT and *Keap1^−/−^*P14 T cells, respectively. Blue circles highlight conventional NRF2 targets from the NFE2L2.V2 gene set (M2870). **(C)** Relative expression levels of *Nqo1* and *Ptgir* mRNA (normalized to *Actb*) in WT and *Keap1^−/−^* P14 T cells as determined by qPCR (mean±SEM, n=3). **(D-E)** Relative expression of **(D)** mRNA and **(E)** protein levels for NAD(P)H quinone dehydrogenase 1 (*Nqo1/*NQO1) and *Ptgir/*PTGIR in activated WT and *Keap1^−/−^* P14 cells following CRISPR-Cas9 editing using control (*sgScr)* or *Nfe2l2*-targeting (*sgNfe2l2*) sgRNAs (mean±SEM, n=3). RNA levels were normalized to *Actnb*. Protein levels were normalized relative to ACTN4 levels. **(F)** Data tracks for NRF2 peak enrichment at *Ptgir* (top) and *Glcm* (bottom) gene loci in CD8 T cells expressing a constitutively active form of NRF2 (CA-NRF2) as determined by NRF2 ChIP-seq. \*\*\**P*<0.001; \*\*\*\**P*<0.0001.

We next assessed whether the induction of *Ptgir* in *Keap1^−/−^*CD8 T cells was dependent on NRF2. CRISPR-Cas9 mediated knockout of the NRF2-encoding gene *Nfe2l2* in *Keap1^−/−^* T cells reduced expression of *Ptgir* mRNA (**Fig. 4D**) and PTGIR protein **(Figs. 4E, S6E-H)** to levels observed in control T cells, indicating that PTGIR expression in *Keap1^−/−^* CD8 T cells was dependent on NRF2. To determine whether NRF2 is a direct transcriptional regulator of *Ptgir* expression, we analyzed NRF2 chromatin immunoprecipitation sequencing (ChIP-seq) data reported by a separate study (Zhu et al, manuscript under revision). NRF2 ChIP-seq revealed enrichment of NRF2 binding upstream of the promoter region of the *Ptgir* gene, with enrichment also observed at the promoter of the known NRF2 target gene *Gclm* **(Fig. 4F)**. Collectively these data establish the prostacyclin receptor PTGIR as a direct transcriptional target of NRF2 that is upregulated in *Keap1^−/−^* CD8 T cells.

### NRF2-mediated expression of PTGIR drives CD8 T cell exhaustion

While our scRNAseq data indicate that NRF2 activity is enriched in terminally exhausted CD8 T cells (**Figs. 2B-C, S3B**), PTGIR expression in CD8 T cell subsets is not known. We analyzed publicly-available transcriptional profiles from CD8 T cell subsets responding to *Listeria monocytogenes* infection or CD8^+^ tumor infiltrating lymphocytes (TILs) derived from an autochthonous murine liver cancer model (*24*) and compared *Ptgir* expression versus NRF2 pathway enrichment scores. We found a positive correlation (R^2^=0.92) of NRF2 enrichment score versus *Ptgir* mRNA expression across CD8 T cell subsets, with the highest *Ptgir* expression found in TIL isolated from established tumors **(Fig. 5A)**. These data indicate a strong correlation between NRF2 activity and *Ptgir* expression in CD8 TIL.

**Fig. 5.**
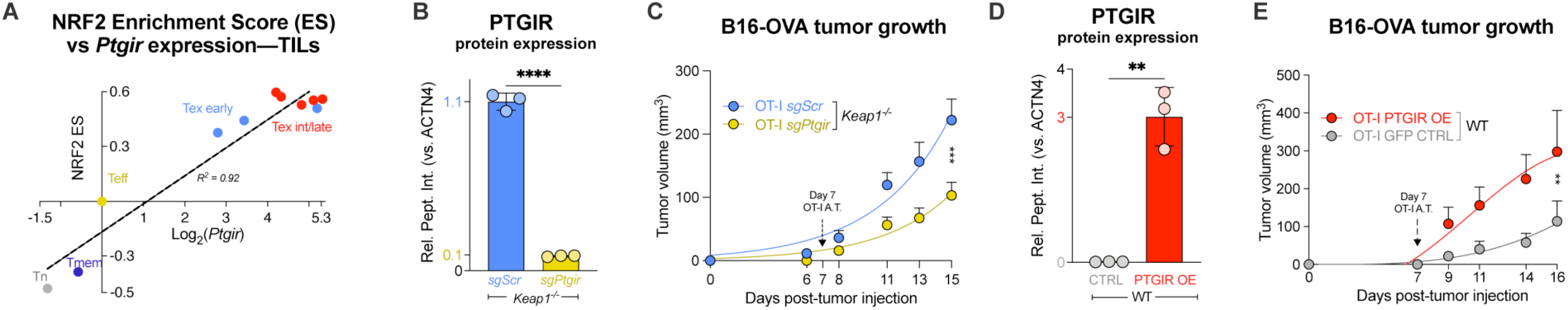
PTGIR is enriched in CD8 TIL and its expression correlates with poor tumor control. **(A)** Correlation between NRF2 activity and *Ptgir* expression in CD8 T cell subsets. Relative *Ptgir* mRNA expression (Log_2_ fold-change) versus NRF2 pathway enrichment scores (NFE2L2.V2) in CD8 T cells from *Listeria monocytogenes* infection (Tn, Teff, and Tmem) or CD8 TIL isolated from early (<21 days) versus late (>21 days) stage liver tumors (data from (*24*)). Expression values were normalized to Teff cells. Linear regression indicates goodness of fit (R^2^=0.92). **(B)** Relative PTGIR protein levels (normalized to ACTN4) in *Keap1^−/−^*CD8 T cells following CRISPR/Cas9 gene editing using sgRNAs targeting *Ptgir* (*sgPtgir*) or non-targeting control (*sgScr*) (mean±SEM, n=3). **(C)** B16-OVA melanoma tumor growth in mice that received adoptive transfer of control (*sgScr*) or PTGIR-deleted (*sgPtgir*) *Keap1^−/−^* OT-I CD8 T cells at 7 dpti (mean±SEM, n=15-17). **(D)** Relative PTGIR protein expression (normalized to ACTN4) in WT OT-I T cells following transduction with empty vector (CTRL) or PTGIR-expressing (PTGIR OE) retroviral vector (mean±SEM, n=3). **(E)** B16-OVA melanoma tumor growth in mice that received adoptive transfer of control (CTRL) or PTGIR-expressing (PTGIR OE) WT OT-I cells at 7 dpti (mean±SEM, n=8). \*\**P*<0.01, \*\*\**P*<0.001, \*\*\*\**P*<0.0001.

To examine the functional role of PTGIR in CD8 T cell responses to cancer, we silenced *Ptgir* in *Keap1^−/−^* OT-I T cells via CRISPR-Cas9 gene editing using guide RNAs targeting *Ptgir* (*sgPtgir*) **(Fig. 5B, Table S2)**. Upon adoptive transfer of PTGIR-ablated *Keap1^−/−^* OT-I T cells into mice bearing B16-OVA tumors, we observed a significant reduction in tumor growth compared to animals that received control (*sgScr*) *Keap1^−/−^* OT-I T cells **(Fig. 5C).** As a complimentary approach, we engineered control (WT) OT-I CD8 T cells to overexpress PTGIR via retroviral-mediated gene transfer (**Fig. 5D**), and B16-OVA tumor bearing mice were administered either control (CTRL) or PTGIR overexpressing (OE) OT-I T cells 7 days post-tumor implantation. Tumor growth was accelerated in mice that received PTGIR OE OT-I T cells versus control empty vector **(Fig. 5E)**. Together, these data indicate that PTGIR expression—driven by NRF2—negatively regulates tumor control by CD8 T cells.

### PTGIR is a checkpoint for CD8 T cell function

Given that increasing PTGIR expression is sufficient to blunt CD8 T cell anti-tumor responses (**Fig. 5E**), we next assessed whether PTGIR functions as an immune checkpoint to reduce CD8 T cell function. To test this, we assessed the impact of reducing *Ptgir* expression on CD8 T cell responses to chronic LCMV infection. We used short hairpin RNAs (shRNAs) targeting *Ptgir* to silence *Ptgir* expression in control (WT) P14 T cells (CD45.2^+^) **(Table S2)**, adoptively transferred them into congenic (CD45.1^+^) hosts, then challenged the mice with LCMV CL13 infection. Flow cytometry analysis of splenocytes on 8 dpi revealed exhibited a >2-fold increase in P14 CD8 T cell expansion when *Ptgir* was silenced (*shPtgir*) compared to control cells (*shCtrl*) (**Figs. 6A, S7A**). In addition, reducing PTGIR expression in P14 CD8 T cells reversed several features exhaustion following LCMV CL13 infection: *shPtgir*-expressing T cells displayed a lower percentage of cells co-expressing the inhibitory receptors PD-1 and TIM-3 **(Fig. 6B, S7B)**, while PD-1^+^ T cells exhibited increased IFN-γ production (**Fig. 6C**). Moreover, PD-1^+^ T cells expressing *shPtgir* displayed increased polyfunctionality marked by heightened co-expression of IFN-γ with both GZMB (**Fig. 6D**) and TNF-α (**Fig. S7C**).

**Fig. 6.**
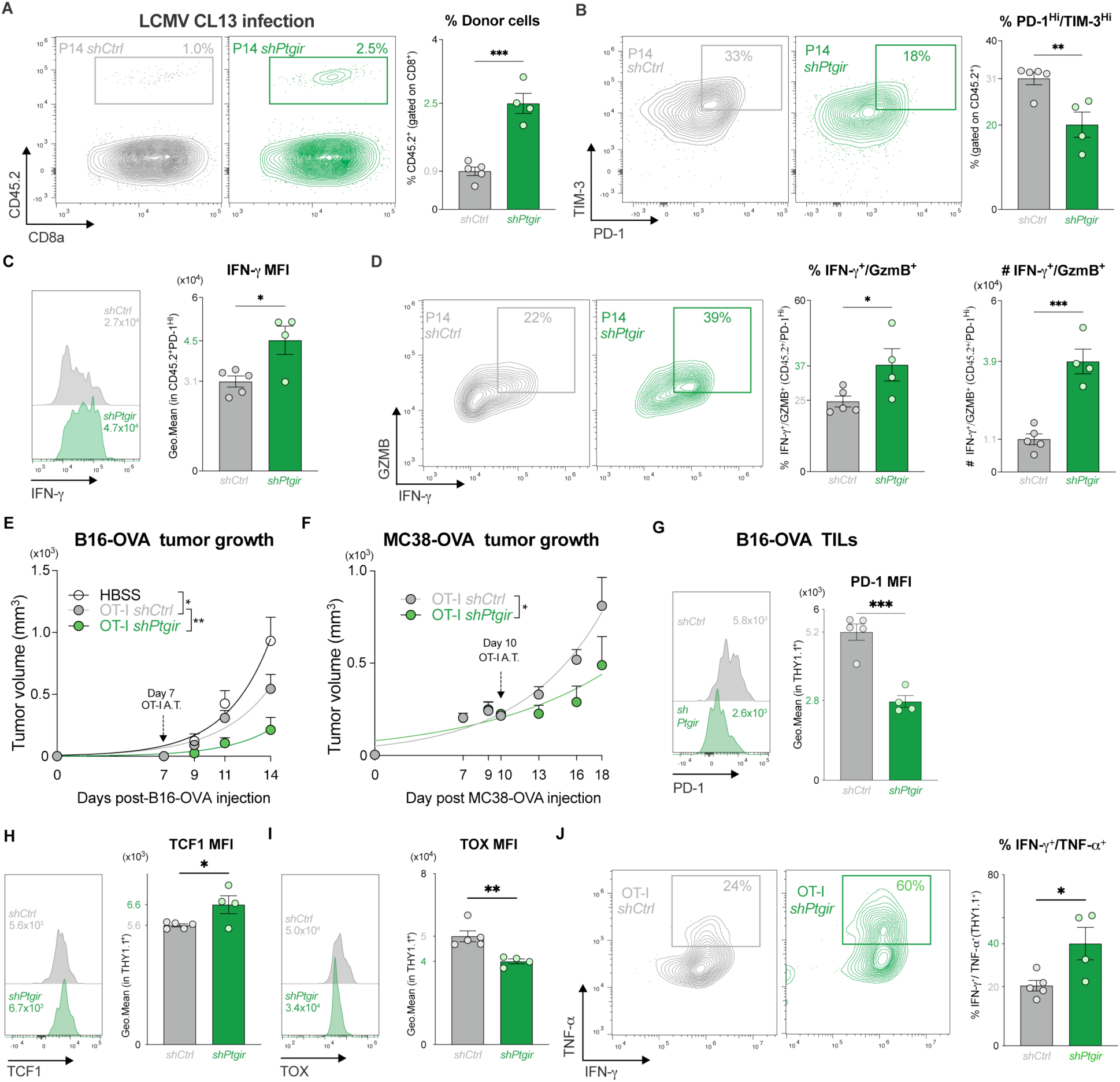
PTGIR is a checkpoint for CD8 T cell function. **(A)** Representative flow cytometry plots of CD8 versus CD45.2 expression on splenocytes harvested from LCMV CL13-infected mice at 8 dpi that received P14 CD8 T cells expressing control (*shCtrl*) or *Ptgir*-targeting *(shPtgir)* shRNAs at −1 dpi. Bar graphs quantify the percentage of donor CD45.2^+^ P14 T cells of total CD8^+^ splenocytes at 8 dpi (mean±SEM, n=4-5/sample). **(B)** Representative flow cytometry plots of PD-1 versus TIM-3 expression by donor CD45.2^+^ P14 T cells expressing control (*shCtrl*) or *Ptgir*-targeting *(shPtgir)* shRNAs at 8 dpi from (A). Bar graph quantifies the percent PD-1^HI^TIM-3^HI^ CD45.2^+^ P14 T cells in the spleen (mean±SEM, n=4-5/sample). **(C)** Histograms and gMFI for IFN-γ expression by PD-1^HI^ P14 CD8 T cells expressing control (*shCtrl*) or *Ptgir*-targeting *(shPtgir)* shRNAs from (A). **(D)** Representative flow cytometry plots of IFN-γ and GZMB expression by PD-1^Hi^ P14 CD8 T cells expressing control (*shCtrl*) or *Ptgir*-targeting *(shPtgir)* shRNAs at 8 dpi as in (A). Bar graph quantifies the percentage and number of polyfunctional (IFN-γ^+^GZMB^+^) P14 T cells in the spleen at 8 dpi (mean±SEM, n=5/sample). **(E-F)** Control of tumor growth by PTGIR-silenced CD8 T cells. Growth of (E) B16-OVA and (F) MC38-OVA tumors following adoptive transfer of *shCtrl*- or *shPtgir*-expressing OT-I CD8 T cells (THY1.1^+^) T cells at 7-10 dpti (mean±SEM, n=4-5). **(G-I)** Representative histograms and bar graph of gMFI for (G) PD-1, (H) TCF1, and (I) TOX expression by *shCtrl*- or *shPtgir*-expressing OT-I TIL from B16-OVA tumors as in (E). **(J)** IFN-γ versus TNF-α expression in *shCtrl*- or *shPtgir*-expressing OT-I TIL from B16-OVA tumors as in (E). Bar graph quantifies the percentage of IFN-γ^+^TNF-α^+^ OT-I TIL from B16-OVA tumors (mean±SEM, n=4-5). \**P*<0.05, \*\**P*<0.01, \*\*\**P*<0.001, \*\*\*\**P*<0.0001.

Finally, we assessed the impact of *Ptgir* knockdown on CD8 T cell-mediated anti-tumor immunity in the context of adoptive cell therapy. Control (WT) OT-I CD8 T cells were transduced with control (*shCtrl*) or *Ptgir*-targeting (*shPtgir*) shRNAs, followed by adoptive transfer into mice bearing B16-OVA tumors. Silencing PTGIR in OT-I T cells resulted in a robust attenuation of B16-OVA tumor growth and increased time to experimental endpoint compared to tumor-bearing mice that received control (*shCtrl*) OT-I T cells **(Fig. 6E, S7D-E)**. Tumor growth results from the B16-OVA model were recapitulated upon adoptive transfer of *shPtgir*-expressing OT-I T cells into mice bearing OVA-expressing MC38 colon carcinoma tumors (**Fig. 6F, S7F-G**). Analysis of TIL from B16-OVA tumors revealed reduced PD-1 expression on PTGIR-deficient OT-I TIL compared to control cells (**Fig. 6G**). Silencing *Ptgir* increased TCF1 expression and decreased TOX expression in OT-I TIL (**Fig. 6H-I**), suggesting a shift in CD8 TIL towards a functional “progenitor exhausted” state, rather than a dysfunctional, terminally exhausted state. Consistent with this, silencing PTGIR expression in OT-I TIL promoted a boost in polyfunctional T cells in B16 tumors, marked by a two-fold increase in the percentage of IFN-γ^+^TNF-α^+^ CD8 T cells compared to control TIL (**Fig. 6J**). Together, these data demonstrate that reducing PTGIR expression in CD8 T cells promotes increased effector function and reduced exhaustion, leading to more robust tumor control. Collectively, our data suggest that PTGIR functions as an immune checkpoint that limits CD8 T cell effector function.

## DISCUSSION

Immune evasion is a critical challenge for cancer therapy (*46*). Current FDA-approved ICI therapies (e.g. Pembrolizumab, Toripalimab) revitalize anti-cancer immune responses in part by preventing immune checkpoint proteins (e.g., PD-1, CTLA-4) from dampening CD8 T cell proliferation and effector function (*47*, *48*). Yet up to 70% of patients treated with ICI therapy relapse or fail to respond to treatment (*49*). Thus, there is a need to better understand the mechanisms that promote CD8 T cell dysfunction and identify additional immune checkpoints that limit CD8 T cell function in the tumor microenvironment (TME). Altered cellular metabolism is a core feature of dysfunctional T cells (*7*, *8*), yet how changes in metabolism impact T cell fate decisions have remained unclear. Here we establish KEAP1-NRF2 signaling as a regulator of T cell exhaustion. Through transcriptional analysis of CD8 TIL and T cells responding to chronic infection, we identified elevated NRF2 activity as a common feature of terminally exhausted (PD-1^+^TIM-3^+^) CD8 T cells. Increasing NRF2 activity in CD8 T cells (via deletion of KEAP1) boosted production of the antioxidant glutathione and lowered cellular ROS levels, consistent with known functions of NRF2 in orchestrating metabolic adaptations that protect cells from oxidative stress (*50*); however, increasing NRF2 activity also promoted the development of Tex^term^ cells in both chronic infection and cancer. Mechanistically, we identified the prostacyclin receptor PTGIR as a direct transcriptional target of NRF2 that promotes terminal exhaustion. Silencing PTGIR restored the anti-tumor function of *Keap1*^−/−^ CD8 T cells and was sufficient to boost both effector function and tumor control by wild type CD8 T cells. Collectively, these data establish the KEAP1-NRF2-PTGIR axis as a regulatory pathway for CD8 T cell exhaustion. We propose that KEAP1 functions as metabolic sensor linking oxidative stress to CD8 T cell function, and that inhibiting NRF2-dependent PTGIR signaling represents a novel strategy for bolstering anti-tumor T cell responses, either alone or in combination with existing ICI therapies.

Glutathione synthesis is critical for promoting T cell activation and protection against ROS (*19*), and ROS has been shown to promote CD8 T cell terminal exhaustion (*18*). Yet paradoxically our data establish that robust activation of NRF2-dependent transcription promotes CD8 T cell dysfunction in vivo in a cell-intrinsic manner despite both enhancing glutathione production and lowering ROS levels. One of the key conclusions of our study is that NRF2 changes the responsiveness of CD8 T cells to cell-extrinsic environmental cues that impact T cell function, notably signals from the prostacyclin receptor PTGIR. Our data establish the *Ptgir* gene as a direct NRF2 target, and that increasing PTGIR levels in CD8 T cells, either through ectopic expression of PTGIR or KEAP1 loss, is sufficient to dampen tumor control by CD8 T cells. PTGIR (and NRF2)-mediated CD8 T cell exhaustion is only observed in vivo and is likely reliant on prostacyclin release from other cell types in the TME. Notably, prostacyclin levels (as inferred by 6-keto-PGF1α) are elevated in many advanced stage tumors such as pancreatic adenocarcinoma (PDAC) compared to normal control tissues (*51*). Furthermore, scRNA-seq of murine and human PDAC tumors indicate that fibroblasts express prostacyclin synthase (*Ptgis/PTGIS*) mRNA and may be the dominant source of prostacyclin in the TME (*51*). The immunosuppressive function of PTGIR may not be restricted to CD8 T cells, as FOXP3 expression and IL-10 secretion by CD4^+^ regulatory T cells (Tregs) is also regulated by prostacyclin-PTGIR signaling (*52*).

Our findings add to a growing literature identifying eicosanoid lipids as negative regulators of CD8 T cell function in vivo. Interaction of prostaglandin E2 (PGE_2_) with its receptors EP2/EP4 has recently been shown to promote CD8 T cell dysfunction in response to chronic antigen stimulation in vivo (*53–55*). During chronic LCMV infection in mice, Chen et al. demonstrated that upregulation of EP2/EP4 receptors on CD8 T cells dampens their survival and cytokine (i.e., IFN-γ, TNF-α) release. Conversely, genetic deletion of EP2/EP4 receptors in CD8 T cells, or pharmacological inhibition of PGE_2_ production via cyclooxygenase (COX1/2) inhibition in combination with anti-PD-L1 treatment, can enhance viral control (*53*). Additionally, two recent manuscripts demonstrated a role for the PGE_2_-EP2/EP4 signaling axis in CD8 TIL dysfunction using BRAF^V600E^ melanoma and MC38 colorectal cancer lines (*54*, *55*). Deletion of the EP2/EP4 receptors on CD8 T cells significantly attenuated tumor growth (*54*), similar to our results when PTGIR expression is reduced in CD8 TIL. Collectively, these data suggest that eicosanoid lipids provide immunosuppressive cues that tune CD8 T cell function in response to conditions in the local environment.

Terminally exhausted CD8 T cells are not inert, as they maintain low levels of cytokine production and limit pathological inflammation during chronic infection (*2*). Accordingly, highly active CD8 T cell responses to viral or cancer antigens can cause excessive damage and even death to the host. For example, intermediate titers of CL13 LCMV (2 ×10^5^ PFU) induce moderate increases in PD-1 expression and robust cytokine production by CD8 T cells, causing the most severe immune pathology, with >75% of infected hosts dying (*56*). Thus, rapidly transitioning to a terminally exhausted state can ultimately be protective to the host. In this light, the induction of terminal exhaustion by NRF2 can be viewed as protective to both T cells and the host. NRF2 activation simultaneously enhances CD8 T cell survival through glutathione-dependent ROS detoxification but at the same time limits T cell-mediated tissue damage by dampening effector function and proliferation. Thus, NRF2-dependent PTGIR expression effectively tunes CD8 T cell responses to the physiologic environment (i.e., tissue prostacyclin levels) while maintaining long-term T cell survival and health of the host.

Our data demonstrating that genetic silencing of PTGIR can enhance the anti-tumor responses of CD8 T cells in vivo highlight the potential of targeting prostacyclin-PTGIR signaling for cancer immunotherapy. Potential applications include the development of blocking antibodies against PTGIR, similar to PD-1/PDL-1 blockade, or ablating PTGIR expression in chimeric-antigen receptor (CAR) T cells for adoptive cell therapy (*57*). Reducing prostacyclin levels in the TME via small molecule inhibitors against prostacyclin synthase (PTGIS) may also antagonize NRF2-dependent T cell dysfunction in viral infection or tumor settings. Defining NRF2 activity and *Ptgir* expression in human TIL will help identify CD8 T subsets receptive to disruption of prostacyclin-PTGIR signaling to ultimately bolster anti-cancer responses. In summary, our data highlight a central role for KEAP1-NRF2 signaling in regulating CD8 T cell fate between effector and exhausted states and highlight a novel role for prostacyclin-PTGIR signaling in the rewiring of CD8 T cell effector function.

## MATERIALS AND METHODS

### Study Design

The objective of this study was to determine how hyperactivation of NRF2 transcriptional targets impacts CD8 T cell exhaustion, and how NRF2-mediated *Ptgir* expression drives the exhausted state. Our RNA-seq meta-analysis defined NRF2 as the most hyperactive transcription factor in CD8 Tex versus effector cells. To investigate the role of NRF2 in CD8 Tex cells, we developed and utilized mouse models with *Keap1* T cell knockout according to approved VAI IACUC-approved protocols and ethical animal treatment. Baseline immunophenotyping revealed no sex-specific differences in examined immune cell types. All in vivo experiments were conducted with at least 4-17 mice and were reproduced 2-3 times. Upon LCMV CL13 (chronic strain) infection, CD8 T cells from *CD4^Cre^ Keap1^fl/fl^ (*termed *Keap1^−/−^*) mice or adoptively transferred LCMV (gp_33-41_)-specific P14-*Keap1^−/−^* T cells were carefully assessed via flow cytometry or viral plaque assays. *Keap1^−/−^* (versus WT) mice were inoculated with B16-OVA melanoma or MC38-OVA colorectal cancer. Alternatively, tumor-bearing mice adoptively transferred with OVA-peptide specific OT-I *Keap1^−/−^*cells were measured for tumor outgrowth. Mechanistic experiments involving *Nfe2l2* (NRF2-encoding) and *Ptgir* knockout were initiated via electroporation-based CRISPR-Cas9 technology, and *Ptgir* knockdown and overexpression experiments were conducted using retroviral transduction according to VAI-approved protocols. ^13^C Stable isotope labeling (SIL) experiments of *Nfe2l2* knockout cells versus control (both in *Keap1^−/−^* background) were conducted under physiologic-like conditions. Fractional enrichment and abundance of M+N isotopologues were analyzed via mass spectrometry. Adoptive transfer experiments with P14 (LCMV infection) or OT-I (B16-OVA/MC38-OVA-targeting) cells bearing *Ptgir* knockdown/knockout were analyzed via flow cytometry of spleens and TIL, respectively.

### RNA-seq meta-analysis

Raw sequences from RNA-seq of CD8^+^ T cells from three previously published studies (GEO accessions: GSE89307, GSE84820, and GSE86881) were downloaded and analysis was conducted as described in the methods section by Luda and Longo et al. (*13*), with the following exceptions: Differential gene expression analyses of all Tex (early, intermediate, late) cells versus Teff cells were conducted on raw counts using DESeq2, with a covariate to adjust for batch, and Benjamini-Hochberg adjusted p-values to maintain a 5% false discovery rate.

### Experimental mouse models

C57BL/6J, CD90.1 (Thy1.1^+^), B6.SJL-*Ptprc^a^ Pepc^b^*/BoyJ (CD45.1^+^), Tg(TcraTcrb)1100Mjb (OT-I), B6.Cg-Tg(Cd4-cre)1Cwi/BfluJ (*Cd4^Cre^*), and B6(Cg)-*Keap1^tm1.1Sbis^*/J (*Keap1^flox/flox^, or Keap1^fl/fl^*) mice were purchased from Taconic Biosciences. Under Van Andel Institute (VAI) Institutional Animal Care and Use Committee (IACUC)-approved protocols, *Keap1^fl/fl^* mice were crossed to *Cd4^Cre^* mice, followed by subsequent breeding of *Keap1^fl/+^* heterozygotes to generate a stable *CD4^Cre^ Keap1^fl/fl^* mouse colony. *CD4^Cre^ Keap1^fl/fl^* were crossed to P14 or OT-I mice followed by subsequent breeding of heterozygotes to generate stable *CD4^Cre^ Keap1^fl/fl^* P14 or *CD4^Cre^ Keap1^fl/fl^* OT-I colonies. Mice were maintained under specific pathogen-free conditions at VAI under approved protocols. Genotyping was performed using DNA extracted from tail or ear biopsies. Experiments were performed using either male or female mice between 8 and 16 weeks of age weighing between 18-25g.

### RNA isolation, sequencing, and qPCR analysis

Total RNA was isolated from murine T cells via RNeasy Kit (#74104, Qiagen) with DNase digestion (#79254, Qiagen) following the manufacturers’ instructions. For quantitative PCR (qPCR) analysis, total RNA was reverse transcribed using a High-Capacity cDNA Reverse Transcriptase kit (#4368814, Life Technologies) and qPCR performed using universal SYBR green supermix (#1725150, Bio-Rad). Results were normalized to *Actb* mRNA levels and control conditions using standard ddCt methods. A list of qPCR primers used in this study can be found in Table S2. RNA preparation and library construction for RNAseq was conducted by the VAI Genomics Core as previously described (*58*). Libraries were sequenced using a NovaSeq 6000 (Illumina) using 50 bp paired-end sequencing (5×10^7^ reads/sample). Gene-set enrichment analysis (GSEA) of RNA-seq data was conducted using the gage function and non-parametric Kolmogorov–Smirnov test from the GAGE (version 2.22.0) R Bioconductor package(*59*). RNA-seq data files are available at NCBI GEO (accession: GSE244465).

### Immunoblotting

Cells were lysed in RIPA or modified Laemmli lysis buffer (240 mM Tris/HCl pH 6.8, 40% glycerol, 8% SDS, 5% 2-ME) supplemented with protease and phosphatase inhibitor cocktails (#4693116001, #PHOSS-RO, Roche). A Pierce BCA Protein Assay Kit (#23225, Thermo Fisher) was used to quantify protein from whole cell lysates. Equal amounts of protein were diluted in Laemmli sample buffer, boiled for 5 min, and resolved by SDS-PAGE on a 10% gel. Proteins were transferred onto PVDF or nitrocellulose membranes. Membranes were blocked for 1 h in 5% non-fat milk in 1X TBST at room temperature and incubated with primary antibodies against KEAP1 (#8047, Cell Signaling), NQO1 (#11451-1-AP, Proteintech), HMOX1(#66743-1-ig, Proteintech) and β-ACTIN (#4967, Cell Signaling) overnight at 4°C. Membranes were washed three times for 5 min with 1X TBST, and then incubated for 1 h at room temperature with the corresponding anti-rabbit IgG (#7074, Cell Signaling) or anti-mouse IgG (#7076, Cell Signaling) HRP-conjugated secondary antibody diluted in 5% non-fat milk. Membranes were washed three times with 1X TBST and then developed using ECL solution (#P132106, ThermoFisher).

### Flow cytometry

Mice were sacrificed and spleens were obtained—or blood was obtained via submental bleed of live mice—and lymphocytes were isolated using red blood cell (RBC) lysis buffer containing 0.15 M NH_4_Cl, 10 mM KHCO_3_, and 0.1 mM EDTA, followed by neutralization with 3 volumes of TCM. Lymphocyte suspensions were surface stained with a cocktail of fluorescently-labeled antibodies: GP33 APC tetramer (custom order, Baylor College of Medicine), KLRG1 AF532 (#58-5893-80, Clone 2F1, eBioscience), CX3CR1 BB700 (#149036, Clone SA011F11), CD8A BUV395 (#563786, Clone 53-6.7, BD Biosciences), CD44 BV650 (#103049, Clone 1M7, Biolegend), CD127 BV785 (#135037, Clone A7R34, Biolegend), TIM-3 BV711 (#119727, Clone RMT3-23, Biolegend), PD-1 BUV737 (#749306, Clone RMPI-30, BD Biosciences) IFN-γ FITC (#17-7311-82, Clone XMG1.2, eBioscience), TNF-α PE-Cy7 (#35-7321-83, Clone MP6-X1222, eBioscience), TOX PE (#130-120-785, Clone REA473, Miltenyi), TCF1 BV421 (#566692, Clone S33-966 RUO, BD Biosciences), Granzyme-B PE/Dazzle594 (#372216, Clone QA16A02, Biolegend), TBET BV605 (#644817, Clone 4B10, Biolegend), CD45.2 BUV805 (#741957, Clone 104, BD Biosciences), Thy1.1 APC (#17-0900-82, Clone HIS51, eBioscience). Cell viability was assessed by using Fixable Viability Dye (FVD) eFluor 506 (#65-0866-14, Invitrogen) or FVD eFluor 780 (#65-0865-18, Invitrogen) according to the manufacturer’s protocols. To assess cytokine production, splenic preps were incubated (37°C, 5% CO_2_ incubator) in TCM in the presence of 1μg/mL GP33_33-41_(KAVYNFATM) peptide (#AS-61669, Anaspec) for 4 h and with (1:1500 dilution) GolgiStop protein transport inhibitor (#554724, BD Biosciences) added for the last 2 h of stimulation prior surface staining. After re-stimulation, cells were surface stained, fixed, and permeabilized using FoxP3/Transcription Factor Staining Buffer Set (#00-5523-00, ThermoFisher), followed by intracellular staining using fluorescently labeled antibodies. Flow cytometry was performed on Cytoflex (Beckman Coulter) or Aurora Cytek cytometers and cell sorting on Astrios (Beckman Coulter) or BD FACSAria Fusion cell sorters. Flow cytometry analysis was performed using FlowJo software (Tree Star).

### Lymphocytic choriomeningitis virus (LCMV) preparation and infection

For preparing viral particles, 10×10^6^ BHK21 cells (#CCL-10, ATCC) in a T-175 cm^2^ flask were infected with either LCMV CL13 or Armstrong virus stocks at a multiplicity of infection (MOI) of 0.1 in 5 ml BHK21 media (see formulation below), and incubated at 37 °C, 5% CO_2_ for 1 hour, rocking every 15 minutes. The supernatant was then removed and replaced with 15 ml of fresh media. Cells were next incubated for 48 hrs at 37°C, 5% CO_2_ and media were collected and centrifuged at 500 RCF for 10 minutes at 4°C. The supernatant was then aliquoted into fresh tubes, and frozen at −80°C until further use following quantitation of plaque forming units (PFU). BHK21 media: 500 mL Dulbecco’s Modified Eagle Medium (DMEM, #319-015, Wisent), 10% (v/v) FBS (#090-150, Wisent), 1% (v/v) of 200 mM L-Glutamine (#25030164, Gibco), 2.8% (v/v) 20% glucose (#A2494001, ThermoFisher), 7% Tryptose phosphate broth (#18050039, ThermoFisher), 1% (v/v) Sodium Pyruvate (#11360070, ThermoFisher), and 1% (v/v) penicillin/streptomycin (#15070063, Gibco).

LCMV Armstrong and LCMV CL13 were diluted to 2×10^5^ and 2×10^6^ PFU, respectively, in 1X D-PBS (#311-425, Wisent) and administered intraperitoneally (Armstrong) or via tail vein (CL13) into host mice.

### LCMV plaque assays

Upon sacrificing, a single mouse kidney was weighed and placed in a 2 mL safelock microcentrifuge tube (#05-402-12, Eppendorf) containing 1 mL kidney extraction buffer: HBSS (#311-512-CL, Wisent), 2% (v/v) FBS, and 1% (v/v) pen./strep. Next, a stainless-steel bead (#69989, Qiagen) was added to each tube and homogenized at 30 rev/s for 3 minutes for 2 rounds using a tissue homogenizer/lyser (#9003240, Qiagen). Kidney extracts were centrifuged at 17,000 RCF for 10 minutes and the supernatant was transferred to a 1.5 mL microcentrifuge tube and kept over ice for 1-2h. Kidney extracts were serially diluted 1:10, 1:33, 1:100, and 1:333 to a final volume of 300 μL in virus dilution media containing: DMEM, +2% v/v FBS. Next 200 μL of kidney extracts diluted in the above media were added to a well within a 24-well plate and 1.4 x 10^5^ MC57G cells (#CRL-2295, ATCC) in 200 μL DMEM containing 10% (v/v) FBS, 1% (v/v) pen./strep., 1% (v/v) of 200 mM L-Glutamine, and 0.1% v/v 2-mercaptoethanol (#21985023, ThermoFisher) were added over top. 24-well plates were gently tapped to ensure an even distribution of cells and incubated at 37°C, 5% CO_2_ for 3-6 hours, or until MC57G cells adhered to the plate in a uniform monolayer. 200 μL of pre-warmed 2X DMEM/2% methylcellulose was added to each well, and incubated at 37°C, 5% CO_2_ for 48h. Once cells reached confluence, media were aspirated, and cells were washed once with 200 μL of HBSS. Next, 200 μL of 10% formalin (#HT501128, Sigma-Aldrich) diluted to 4% in 1X PBS was added overtop of cells for 20 minutes, followed by two washes with 200 μL PBS. After, 200 μL 0.5% Triton-X (#X-100, Sigma-Aldrich) in HBSS was added overtop for 20 minutes, followed by one wash with 200 μL PBS. For primary antibody labelling, PBS was aspirated from each well and 200 μL of 10% FBS in PBS was added overtop of cells for 60-90 minutes at room temperature (RT), followed by 1 wash with 200 μL PBS. 200 μL of VL-4 rat anti-LCMV monoclonal antibody (diluted 1:10 in PBS+1% w/v FBS, #B10106, Bio X Cell) was added overtop for 30-60 minutes at RT, followed by three washes with 1X PBS. Next, 200 μL of peroxidase conjugated goat anti-rat secondary antibody (#112-035-003, Jackson ImmunoResearch), diluted 1:200 in PBS+1% (w/v) FBS was added overtop of cells and incubated for 30-60 minutes at RT, followed by 3 washes with PBS. To obtain plaques, wells were incubated (10-30 minutes) with 200 μL of aqueous color reaction buffer: 0.1M Citric acid (#C0759, Sigma-Aldrich), 0.2M Na_2_HPO_4_*2H_2_O (#71643, Sigma-Aldrich), o-phenylenediamine dihydrochloride (OPD) (#P3804, Sigma-Aldrich), and 100 μL 30% (w/w) hydrogen peroxide (#H1009, Sigma-Aldrich). Colored plaques were then counted and normalized to dilution factor and kidney weight to obtain a plaque forming unit (PFU) value.

### CD8^+^ T cell purification and culture

P14^+/-^, OT-I^+/-^ or total CD8^+^ T cells were purified from spleen and peripheral lymph nodes by negative selection using the EasySep^TM^ Mouse CD8^+^ T cell isolation kit (#19753, StemCell Technologies). For in vitro experiments, isolated cells were first stained with 0.5 μg/mL propidium iodide (Invitrogen) and FITC-CD8A (clone 53-6.7, eBioscience) and assessed for purity (> 95%) by fluorescence-activated cell sorting (FACS). In vitro-activated CD8^+^ Teff cells were generated by stimulating naïve CD8^+^ T cells (1 x 10^6^ cells/mL) with plate-bound anti-CD3ε (clone 145-2C11) and anti-CD28 (clone 37.51) antibodies (eBioscience) for 2 days. Cells were cultured in T cell media—hereafter referred to as TCM—containing Iscove’s Modified Dulbecco’s Medium supplemented with 1% v/v pen./strep., 50 μM of 2-mercaptoethanol, 6 mM L-glutamine, and 10% FBS.

### Cell lines

293T (CRL-3216, ATCC) B16-OVA (SCC420, Sigma), and MC38-OVA (kindly provided by Dr. John Stagg) cells were cultured in DMEM supplemented with 10% v/v FBS, 1% v/v pen.strep, and L-glutamine at a final concentration of 6 mM. All cells were cultured at 37°C in a humidified 5% CO_2_ incubator.

### Retrovirus and Lentivirus production and T cell transduction

For retrovirus production corresponding to PTGIR knockdown in CD8 T cells, 293T cells were first co-transfected with pCL-Eco and either pLMPd-Ametrine shFF (control vector, *shCtr*l) or pLMPd-Ametrine *shPtgir* (**Table S2**)—developed according to the cloning pipeline by Dow et al. (*60*)— using Lipofectamine 2000 transfection reagent (#11668019, Invitrogen). Viral supernatants were harvested 48 and 72 h post-transfection, pooled, and concentrated using Lenti-X Concentrator (#631232, Takara) according to the manufacturer’s protocol. The concentrated retrovirus was added to 24 h-activated T cells with 8 ng/mL polybrene (#TR-1003, Sigma-Aldrich), 200 U/mL IL-2 (#212-12, Thermo Fisher), and 20 mM HEPES, and the cells were centrifuged at 1180 RCF, 30°C for 90 min. Ametrine-positive cells were sorted by FACS 72 h post-transduction, cultured for 48h in T cell media, phenotypically validated for PTGIR knockdown, and adoptively transferred into LCMV CL13-infected or tumor-bearing hosts. For retrovirus production corresponding to PTGIR overexpression in CD8 T cells, 293T cells were first co-transfected with pCL-Eco and either MIGR1 (GFP control vector, kindly provided by Warren Pear, #27490 Addgene), or MIGR1-PTGIR (custom PTGIR overexpressing vector VectorBuilder), using Lipofectamine 2000 transfection reagent and subsequent procedures as mentioned above, with the following exception: 72h post-transduction GFP-positive cells were flow sorted, cultured for 48h in T cell media, phenotypically validated for PTGIR overexpression, and adoptively transferred into tumor-bearing hosts.

### B16-OVA and MC38-OVA tumor model

8–12 week-old male or female *Cd4^Cre^ Keap1^fl/fl^*(*Keap1*^−/−^) mice or *Keap1^fl/fl^* (WT) control littermates were injected with 5 x 10^5^ MC38-OVA cells or 5 x 10^5^ B16-OVA subcutaneously in the right abdominal flank. Once palpable tumors were present, tumor measurements were obtained every 2-3 days using a caliper. Mice were euthanized as they reached humane endpoints, which included a maximum tumor volume of ≥1500 mm^3^. For adoptive transfer experiments, mice were inoculated with 5 ×10^5^ MC38-OVA or B16-OVA cancer cells and on day 7 post inoculation, and when palpable tumors were present, 1×10^6^ OT-I T cells were adoptively transferred into tumor-bearing hosts.

### TIL isolation

On day 7 post adoptive transfer (Day 14-15 post cancer inoculation) mice were sacrificed and respective B16-OVA tumors were measured and excised using a scalpel. In a 6-well plate, tumors were homogenized and passaged three times through a 100-micron filter, followed by a 40-micron filter. Tumor extracts were incubated with RBC lysis buffer for 1 minute at room temperature, and three volumes of TCM was added overtop to neutralize the RBC lysis reaction. TIL were then centrifuged at 500 RCF for 5 minutes, media were aspirated, and resuspended in 0.5-1 mL of TCM and used for downstream flow cytometry experiments.

### CRISPR-Cas9-mediated gene editing

Electroporation of WT and *Keap1^−/−^* P14 and OT-I CD8 T cells was adapted from the method developed by Nussing et al. (*61*). First, antigen-specific (P14 or OT-I) CD8 T cells were isolated from respective mice, assessed for purity (>95%) and activated for 48h with plate bound antibodies in the same fashion as in the above *CD8+T cell purification and culture* section. After 48h of culture, cells were counted and 2.5 ×10^6^ cells were centrifuged at 500 RCF for 5 minutes and washed twice with PBS. During the washes, 40 pmol of SpCas9 2NLS (Synthego) was complexed with 300 pmol of sgRNA (Synthego, see Table S2) in a total volume of 5 μL for 10 minutes. After the final PBS wash, cells were resuspended in 20 μL of P3 buffer mix (#V4XP-3032, Lonza) according to the manufacturer’s protocol and mixed with the 5 μL of pre-incubated Cas9/sgRNA ribonucleoprotein (RNP) complex. The cell/RNP mixture was then immediately added to a 16 well nucleofector strip and electroporated using program DN-100 of 4D-nucleofector X system (#AAF-1003X, Lonza). Following electroporation, the cell/RNP mixture was topped with 130 μL P/S free TCM, then placed in a final volume of (P/S-free) 2.5 mL TCM containing 50 U/mL IL-2 in a 12 well plate, and incubated at 37 °C (5% CO_2_) overnight. The following day, media were changed to fresh TCM containing P/S and 50 U/mL IL-2. Cells were moved to a 6 well plate and diluted to 0.4 x 10^6^ cells/mL and cultured for 48h. Knockout efficiency was assessed via qPCR for NRF2 target genes or via proteomics for PTGIR.

### ^13^C-Glutamine Stable Isotope Labeling (SIL)

Following initial T cell isolation, activation and expansion in TCM for 4 days, WT or *Keap1^−/−^* CD8 T cells were traced according to the method developed by Kaymak and Luda et al. (*12*). For ^13^C tracing, cells were resuspended in 0.8 mL Van Andel Institute Media (VIM), using specialized IMDM (#319-267-CL, Wisent) as a base solution containing 10% (v/v) dialyzed FBS, 1% (v/v) P/S, and 0.1% (v/v) β-Mercaptoethanol, and ^12^C isotopologues of 5 mM glucose, 0.1 mM glycine, 0.2 mM alanine, 0.01 mM aspartate, 0.02 mM glutamate, 0.04 mM methionine, 0.1 mM serine, and 0.1 mM threonine. For ^13^C-[U]-Glutamine (#CLM-1822, Cambridge Isotope Laboratories) tracing, a 5X 0.2 mL of base solution containing 5 mM ^13^C-[U]-Glutamine (0.5 mM final concentration)—containing all ^12^C metabolites at aforementioned concentrations with 250 U/mL IL-2—was added overtop of 0.8 mL cell suspension and incubated for 4h under standard culture conditions. For unlabeled controls, a separate set of cells were incubated with all aforementioned ^12^C metabolites, but with ^12^C glutamine rather than ^13^C glutamine. After 4h incubation, cell solutions were placed into 1.5 mL collection tubes, and centrifuged at 600 RCF for 4 minutes. Media were collected from each sample and frozen over dry ice for 5 minutes and stored at −80°C. Cell pellets were then washed twice with 0.9% w/v saline (NaCl) by centrifuging cell pellets at 600 RCF for 4 minutes. At the last wash, virtually all the saline was removed, and cell pellets were frozen over dry ice for 5 minutes then placed in −80°C freezer for downstream metabolomics analysis.

### ^13^C metabolomics analysis

Metabolites were extracted in ice-cold acetonitrile:methanol:water (40%:40%:20% v/v) at a concentration of 2×10^6^ cells/mL. Crude extracts were vortexed for 30s, sonicated in a water bath sonicator for 5 minutes, and incubated on wet ice for 1 hour. Extracts were precipitated by centrifugation at 14,000 RCF for 10 min at 4°C, and the 760µL of the soluble supernatant (1.52×10^6^cell equivalents), were dried in a vacuum evaporator. Dried extracts were resuspended in 50µL of LCMS grade water, resulting in a final concentration of 3.0×10^7^ cell equivalents/mL. Stable isotope tracing metabolomics data were collected on an Orbitrap Exploris 240 mass spectrometer (Thermo) using a tributylamine ion-paired reversed phase chromatography. The column was a ZORBAX Rapid Resolution HD (2.1 × 150 mm, 1.8 µm; 759700–902, Agilent). Mobile phase A was 3% methanol, and mobile phase B was 100% methanol. Both mobile phases contained 10 mM tributylamine (90780, SigmaAldrich, St Louis, MO, USA), 15 mM acetic acid and 2.5 µM medronic acid (5191–4506, Agilent Technologies, Santa Clara, CA, USA). The LC gradient was: 0–2.5 min 0% B, 2.5–7.5 min ramp to 20% B, 7.5–13 min ramp to 45% B, 13–20 min ramp to 99% B, 20–24 min hold at 99% B. Flow rate was 0.25 mL/min, and the column compartment was heated to 35°C. The column was then backflushed with 100% acetonitrile for 4 min (ramp from 0.25 to 0.8 mL/min in 1.5 min) and re-equilibrated with mobile phase A for 5 min at 0.4 mL/min. The H-ESI source was operated at spray voltage of −2500 V, sheath gas: 60 a.u., aux gas: 19 a.u., sweep gas: 1 a.u., ion transfer tube: 300°C, vaporizer: 250°C. Full scan MS1 data was collected from 70 to 800 m/z at mass resolution of 240,000 FWHM with RF lens at 35%, and standard automatic gain control (AGC). ddMS2 data were collected on unlabeled control samples using HCD fragmentation HCD at stepped collision energies of 20, 35, and 50%., with orbitrap detection at 15000 FWHM. Peak picking and integration were completed in Skyline (v23.3) using an in-house curated compound data base from analytical standards. Natural abundance correction was performed using IsoCorrectoR (*62*). Metabolomics analysis was conducted by the VAI Mass Spectrometry Core (<u>RRID:SCR_024903</u>).

### RNA-sequencing analysis

WT and *Keap1^−/−^* CD45.2^+/+^; P14^+^ cells were adoptively transferred into B6.SJL-*Ptprc^a^ Pepc^b^*/BoyJ (CD45.1^+/+^) hosts one day prior to LCMV Armstrong infection, and P14 cells were isolated on 7 dpi using the EasySep Mouse APC Positive Selection Kit II (#17667, STEMcell technologies) with a CD45.2 APC antibody (#17-0454-82, Clone 104, eBioscience), followed by FACS for additional purification (~100% purity). Sorted cells were then processed for RNA-seq by the VAI Pathology and Biorepository Core, and libraries for RNA-seq were prepared by the VAI genomics core. The VAI Bioinformatics and Biostatistics Core conducted differential expression analysis from *Keap1* KO and WT P14 cells (Figs. 4B, S6A-D, Tables S3-4). Specifically, adaptor sequences and low-quality reads of RNA-seq reads were trimmed using Trim Galore (*63*). Trimmed reads were aligned to the mm10 reference genome using STAR (*64*), with parameter “--quantMode GeneCounts”. Differential gene expression (DGE) analyses were conducted using edgeR (*65–67*), based on gene counts generated from STAR. Batch information was included in the edgeR model as a covariate. For DGE, Benjamini-Hochberg adjusted p-values was set to 0.05. ClusterProfiler (v4.0.5;(*68*)) was used to perform gene set enrichment analysis (GSEA) on gene collections C2, C5, C6, C7, NFE2L2.v2 and H. Data are available on NCBI GEO: GSE244465.

### Proteomics sample preparation, LC-MS/MS, and analysis

Cell lysates were extracted and digested using EasyPrep MS Sample Prep Kits (Thermo Fisher Scientific) according to the manufacturer’s instructions. Dried samples were resuspended in 0.1% TFA. Synthetic peptides (GFTQAIAPDS, EMGDLLAFR, DDPVTNLNNAFEVAEK, and LVSIGAEEIVDGNAK) were obtained from Biosynth (Gardner, MA).

For global Proteome Quantitation Analysis, Data-independent acquisition (DIA) analyses were performed on an Orbitrap Eclipse mass spectrometer coupled with the Vanquish Neo LC system (Thermo Fisher Scientific). The FAIMS Pro source was positioned between the nanoESI source and the mass spectrometer. A total of 2 μg of digested peptides were separated on a nano capillary column (20 cm × 75 μm I.D., 365 μm O.D., 1.7 μm C18, CoAnn Technologies, Washington) at a flow rate of 300 nL/min. Mobile phase A was water with 0.1% formic acid, and mobile phase B was 20% water and 80% acetonitrile with 0.1% formic acid. The LC gradient was programmed as follows: 1% B to 24% B over 110 minutes, 85% B over 5 minutes, and 98% B over 5 minutes, resulting in a total gradient length of 120 minutes. For FAIMS, selected compensation voltages (−40V, −55V, −70V) were applied throughout the LC-MS/MS runs. Full MS spectra were collected at a resolution of 120,000 (FWHM), and MS2 spectra at 30,000 (FWHM). Standard automatic gain control (AGC) targets and automatic maximum injection times were used. A precursor range of 380-980 m/z was set for MS2 scans, with a 50 m/z isolation window and a 1 m/z overlap for each scan cycle. A 32% higher-energy collisional dissociation (HCD) collision energy was used for MS2 fragmentation.

To generate a hybrid library for directDIA™ analysis in Spectronaut, pooled samples underwent data-dependent acquisition (DDA) employing 11 distinct FAIMS CV settings ranging from −30 to 80 CV. Full MS spectra were collected at 120,000 resolution (FWHM), and MS2 spectra at 30,000 resolutions (FWHM). The standard AGC target and automatic maximum injection time were selected. Ions with charges of 2–5 were filtered. An isolation window of 1.6 m/z was used with quadrupole isolation mode, and ions were fragmented using HCD with a collision energy of 32%.

For the targeted quantitation, parallel reaction monitoring (PRM) was performed on an Exploris 480 mass spectrometer coupled with the Vanquish Neo LC system (Thermo Fisher Scientific). A total of 2 μg of digested peptides were separated on a nano capillary column (20 cm × 75 μm I.D., 365 μm O.D., 1.7 μm C18, CoAnn Technologies, Washington) at a flow rate of 300 nL/min. Mobile phase A was water with 0.1% formic acid, and mobile phase B was 20% water and 80% acetonitrile with 0.1% formic acid. The LC gradient was as follows: 1% B to 26% B over 51 minutes, 85% B over 5 minutes, and 98% B over 4 minutes, with a total gradient length of 60 minutes. Full MS spectra (m/z 375-1200) were collected at a resolution of 120,000 (FWHM), and MS2 spectra at 30,000 resolutions (FWHM). Standard AGC targets and automatic maximum injection times were used for both full and MS2 scans. A 32% HCD collision energy was used for MS2 fragmentation. All samples were analyzed using a multiplexed PRM method with a scheduled inclusion list containing the target precursor ions. Two unique peptides of PTGIR (GFTQAIAPDS and EMGDLLAFR) were used to measure the relative abundance of PTGIR, while two alpha-actinin4 peptides (DDPVTNLNNAFEVAEK and LVSIGAEEIVDGNAK) served as internal standards.

DIA data were processed in Spectronaut (version 18, Biognosys, Switzerland) using directDIA™ analysis. Data were searched against the *Mus musculus* reference proteome database (Uniprot, Taxon ID: 10090) with the manufacturer’s default parameters. Briefly, trypsin/P was set as the digestion enzyme, allowing for two missed cleavages. Cysteine carbamidomethylation was set as a fixed modification, while methionine oxidation and protein N-terminus acetylation were set as variable modifications. Identification was performed using a 1% q-value cutoff at both the precursor and protein levels. Both peptide precursor and protein false discovery rates (FDR) were controlled at 1%. Ion chromatograms of fragment ions were used for quantification, with the area under the curve between the XIC peak boundaries calculated for each targeted ion. DDA raw files were utilized in Library Extension Runs to enhance proteome coverage to generate a hybrid library. All PRM data analysis and integration were performed using Skyline software. The transitions’ intensity rank order and chromatographic elution were required to match those of a synthetic standard for each measured peptide. All proteomics sample preparation, LC-MS/MS and analysis was conducted by the VAI mass spectrometry core <u>RRID:SCR_024903</u>.

### ChIP-Sequencing

The detail of ChIP-Seq experiment with anti-NRF2 antibody has been described in detail elsewhere (Zhu et al, manuscript under revision). In brief, the truChIP Chromatin Shearing Kit (#PN520154, Covaris) was used to cross-link DNA-protein, isolate and prepare nuclei and shear chromatin to 200-700bp fragments. Immunoprecipitation was performed with an anti-NRF2 polyclonal antibody (#PA5-27882, Invitrogen). ChIPmentation was conducted to generate the ChIP-seq libraries according to a previously published method described in Schmidel et al. (*69*), and ChIP-seq libraries were sequenced with an Illumina NovaSeq 6000.

### Statistical Analysis

Data are presented as mean ± standard deviation (SD) for technical replicates and mean ± standard error of the mean (SEM) for biological replicates. Observations from technical replicate data were reproduced in at least two independent experiments. Statistical analyses were performed using GraphPad Prism software (GraphPad). A two-tailed, unpaired Student’s t-test was used to compare the means between two independent groups, with the exception of co-transfer experiments, where a paired parametric two-tailed t-test was used. Tumor growth curve statistical analysis between groups was conducted via mixed-effects model with exponential (Malthusian) growth non-linear, or third-order polynomial (cubic) fit. Kaplan-Meier curves from tumor-bearing mice were analyzed via Log-rank (Mantel-Cox) test. Lastly, statistical testing of mass isotopologue distribution (MID) upon [U-^13^C-Glutamine] tracing between groups was conducted via multiple unpaired t-tests using the Holm-Šídák method. Statistical significance is indicated in all figures by the following annotations: *, *p* < 0.05; **, *p* < 0.01; ***, *p* < 0.001; ****, *p* < 0.0001; ns, not significant.

## Supplementary materials

### Supplementary Figures

Fig. S1. Validation of *Keap1* deletion in CD8 T cells.

Fig. S2. Immunophenotyping of splenocyte populations from *Keap1^fl/fl^* (WT) and *Cd4^Cre^ Keap1^fl/fl^*(*Keap1^−/−^*) mice.

Fig. S3. scRNA-seq cluster analysis and impact of *Keap1* deletion in T cells on tumor growth and survival.

Fig. S4. Proteomic and metabolic profiling of *Keap1^−/−^* CD8 T cells.

Fig. S5. Immunophenotyping of *Keap1^−/−^ sgScr* and *sgNfe2l2* P14 cells following LCMV CL13 infection.

Fig. S6. Enriched gene sets in *Keap1^−/−^* versus WT P14 cells and mass spectrometry validation of PTGIR peptides.

Fig. S7. *Ptgir* knockdown improves anti-viral CD8 T cell responses and attenuates tumor growth.

### Supplementary Tables

**Table S1: Gene Set Enrichment Analysis (GSEA) of the oncogenic signature gene sets (MSigDB C6 gene set) enriched in Tex versus Teff cell clusters.**

**Table S2: qPCR, shRNA and sgRNA oligonucleotide sequences.**

**Table S3: Differential gene expression analysis from RNA-seq of adoptively transferred *Keap1^−/−^*versus WT P14 cells, isolated 7 days post LCMV Armstrong infection.**

**Table S4: C2, C5, C6, C7, NFE2L2.v2 and H Gene set enrichment analysis (GSEA) from RNA-seq of *Keap1^−/−^*versus WT P14^+^ CD8^+^ T cells upon LCMV Armstrong infection.**

## Acknowledgments

We acknowledge Sarah E. LeBoeuf and Thales Papagiannakopolous for providing sgRNA sequences to target NRF2 and Ralph DeBerardinis for scientific discussions contributing to this manuscript. We thank Jeanie Wedberg, Margene Brewer, Michelle Minard, and Carly Kraus for administrative assistance and support. We thank members of the Van Andel Institute (VAI) Core Facilities (Metabolomics and Bioenergetics, Genomics, Bioinformatics and Biostatistics, Flow Cytometry, Vivarium) for technical assistance.

## Author contributions

Conceptualization: MSD, RGJ, CMK

Methodology: MSD, RGJ, RSD, HL, AJ, EEE, MV, IS

Investigation: MSD, LMD, BMO, SEC, JL

Visualization: MV, DF

Funding acquisition: MSD, RGJ

Project administration: KSW

Supervision: RGJ, CY, TW,

Writing – original draft: MSD, RGJ, SMG, RDS, HL

Writing – review & editing: MSD, RGJ

## Data and materials availability

All data are available in the main text, supporting figures/tables, or data repositories. The RNA-seq data used for the meta-analysis in Figure 1 are available at NCBI GEO: GSE86881, GSE89307, GSE84820. Bulk ex vivo RNA-seq data of WT and *Keap1^−/−^* CD8^+^ P14^+^ corresponding to Figures 4 and S6 are available at NCBI GEO: GSE244465 (token wrklcuaipvsbfkz). Proteomics data corresponding to Figures 3-5 are uploaded onto PRIDE database (PXD052688).

Original western blot images have been deposited at Mendeley and are publicly available as of the date of publication. DOI:

Any additional information required to reanalyze the data reported in this paper is available from the lead contact upon request.

All unique/stable reagents generated in this study will be made available from the Lead Contact with a completed Materials Transfer Agreement.

Additional information and request for resources and reagents should be directed to and will be fulfilled by the lead contact, Russell G. Jones.

## References and Notes

1. L. M. McLane, M. S. Abdel-Hakeem, E. J. Wherry, CD8 T cell exhaustion during chronic viral infection and cancer. Annu. Rev. Immunol. 37, 457–495 (2019).

2. E. J. Wherry, M. Kurachi, Molecular and cellular insights into T cell exhaustion. Nat. Rev. Immunol. 15, 486–499 (2015).

3. D. M. Pardoll, The blockade of immune checkpoints in cancer immunotherapy. Nat. Rev. Cancer 12, 252–264 (2012).

4. G. Morad, B. A. Helmink, P. Sharma, J. A. Wargo, Hallmarks of response, resistance, and toxicity to immune checkpoint blockade. Cell 184, 5309–5337 (2021).

5. P. Sharma, S. Hu-Lieskovan, J. A. Wargo, A. Ribas, Primary, adaptive, and acquired resistance to cancer immunotherapy. Cell 168, 707–723 (2017).

6. D. Hanahan, R. A. Weinberg, Hallmarks of cancer: the next generation. Cell 144, 646–674 (2011).

7. D. G. Roy, I. Kaymak, K. S. Williams, E. H. Ma, R. G. Jones, Immunometabolism in the Tumor Microenvironment. Annu. Rev. Cancer Biol. 5, 137–159 (2021).

8. F. Franco, A. Jaccard, P. Romero, Y.-R. Yu, P.-C. Ho, Metabolic and epigenetic regulation of T-cell exhaustion. Nat Metab 2, 1001–1012 (2020).

9. Z. A. Bacigalupa, M. D. Landis, J. C. Rathmell, Nutrient inputs and social metabolic control of T cell fate. Cell Metab. 36, 10–20 (2024).

10. M. D. Buck, R. T. Sowell, S. M. Kaech, E. L. Pearce, Metabolic Instruction of Immunity. Cell 169, 570–586 (2017).

11. N. M. Chapman, H. Chi, Metabolic adaptation of lymphocytes in immunity and disease. Immunity 55, 14–30 (2022).

12. I. Kaymak, K. M. Luda, L. R. Duimstra, E. H. Ma, J. Longo, M. S. Dahabieh, B. Faubert, B. M. Oswald, M. J. Watson, S. M. Kitchen-Goosen, L. M. DeCamp, S. E. Compton, Z. Fu, R. J. DeBerardinis, K. S. Williams, R. D. Sheldon, R. G. Jones, Carbon source availability drives nutrient utilization in CD8+ T cells. Cell Metab. 34, 1298–1311.e6 (2022).

13. K. M. Luda, J. Longo, S. M. Kitchen-Goosen, L. R. Duimstra, E. H. Ma, M. J. Watson, B. M. Oswald, Z. Fu, Z. Madaj, A. Kupai, B. M. Dickson, L. M. DeCamp, M. S. Dahabieh, S. E. Compton, R. Teis, I. Kaymak, K. H. Lau, D. P. Kelly, P. Puchalska, K. S. Williams, C. M. Krawczyk, D. Lévesque, F.-M. Boisvert, R. D. Sheldon, S. B. Rothbart, P. A. Crawford, R. G. Jones, Ketolysis drives CD8+ T cell effector function through effects on histone acetylation. Immunity 56, 2021–2035.e8 (2023).

14. E. H. Ma, M. S. Dahabieh, L. M. DeCamp, I. Kaymak, S. M. Kitchen-Goosen, B. M. Oswald, J. Longo, D. G. Roy, M. J. Verway, R. M. Johnson, B. Samborska, L. R. Duimstra, C. A. Scullion, M. Steadman, M. Vos, T. P. Roddy, C. M. Krawczyk, K. S. Williams, R. D. Sheldon, R. G. Jones, 13C metabolite tracing reveals glutamine and acetate as critical in vivo fuels for CD8 T cells. Science Advances 10, eadj1431 (2024).

15. B. Bengsch, A. L. Johnson, M. Kurachi, P. M. Odorizzi, K. E. Pauken, J. Attanasio, E. Stelekati, L. M. McLane, M. A. Paley, G. M. Delgoffe, E. J. Wherry, Bioenergetic Insufficiencies Due to Metabolic Alterations Regulated by the Inhibitory Receptor PD-1 Are an Early Driver of CD8+ T Cell Exhaustion. Immunity 45, 358–373 (2016).

16. Y.-R. Yu, H. Imrichova, H. Wang, T. Chao, Z. Xiao, M. Gao, M. Rincon-Restrepo, F. Franco, R. Genolet, W.-C. Cheng, C. Jandus, G. Coukos, Y.-F. Jiang, J. W. Locasale, A. Zippelius, P.-S. Liu, L. Tang, C. Bock, N. Vannini, P.-C. Ho, Disturbed mitochondrial dynamics in CD8+ TILs reinforce T cell exhaustion. Nat. Immunol. 21, 1540–1551 (2020).

17. N. E. Scharping, D. B. Rivadeneira, A. V. Menk, P. D. A. Vignali, B. R. Ford, N. L. Rittenhouse, R. Peralta, Y. Wang, Y. Wang, K. DePeaux, A. C. Poholek, G. M. Delgoffe, Mitochondrial stress induced by continuous stimulation under hypoxia rapidly drives T cell exhaustion. Nat. Immunol. 22, 205–215 (2021).

18. S. A. Vardhana, M. A. Hwee, M. Berisa, D. K. Wells, K. E. Yost, B. King, M. Smith, P. S. Herrera, H. Y. Chang, A. T. Satpathy, M. R. M. van den Brink, J. R. Cross, C. B. Thompson, Impaired mitochondrial oxidative phosphorylation limits the self-renewal of T cells exposed to persistent antigen. Nat. Immunol., doi: 10.1038/s41590-020-0725-2 (2020).

19. T. W. Mak, M. Grusdat, G. S. Duncan, C. Dostert, Y. Nonnenmacher, M. Cox, C. Binsfeld, Z. Hao, A. Brüstle, M. Itsumi, C. Jäger, Y. Chen, O. Pinkenburg, B. Camara, M. Ollert, C. Bindslev-Jensen, V. Vasiliou, C. Gorrini, P. A. Lang, M. Lohoff, I. S. Harris, K. Hiller, D. Brenner, Glutathione Primes T Cell Metabolism for Inflammation. Immunity 46, 675–689 (2017).

20. L. Baird, M. Yamamoto, The molecular mechanisms regulating the KEAP1-NRF2 pathway. Mol. Cell. Biol. 40 (2020).

21. Y. Boie, T. H. Rushmore, A. Darmon-Goodwin, R. Grygorczyk, D. M. Slipetz, K. M. Metters, M. Abramovitz, Cloning and expression of a cDNA for the human prostanoid IP receptor. J. Biol. Chem. 269, 12173–12178 (1994).

22. T. Kobayashi, F. Ushikubi, S. Narumiya, Amino acid residues conferring ligand binding properties of prostaglandin I and prostaglandin D receptors. Identification by site-directed mutagenesis. J. Biol. Chem. 275, 24294–24303 (2000).

23. K. Man, S. S. Gabriel, Y. Liao, R. Gloury, S. Preston, D. C. Henstridge, M. Pellegrini, D. Zehn, F. Berberich-Siebelt, M. A. Febbraio, W. Shi, A. Kallies, Transcription factor IRF4 promotes CD8+ T cell exhaustion and limits the development of memory-like T cells during chronic infection. Immunity 47, 1129–1141.e5 (2017).

24. M. Philip, L. Fairchild, L. Sun, E. L. Horste, S. Camara, M. Shakiba, A. C. Scott, A. Viale, P. Lauer, T. Merghoub, M. D. Hellmann, J. D. Wolchok, C. S. Leslie, A. Schietinger, Chromatin states define tumour-specific T cell dysfunction and reprogramming. Nature 545, 452–456 (2017).

25. B. Daniel, K. E. Yost, S. Hsiung, K. Sandor, Y. Xia, Y. Qi, K. J. Hiam-Galvez, M. Black, C. J Raposo, Q. Shi, S. L. Meier, J. A. Belk, J. R. Giles, E. J. Wherry, H. Y. Chang, T. Egawa, A. T. Satpathy, Divergent clonal differentiation trajectories of T cell exhaustion. Nat. Immunol. 23, 1614–1627 (2022).

26. M. Yamamoto, T. W. Kensler, H. Motohashi, The KEAP1-NRF2 system: A thiol-based sensor-effector apparatus for maintaining redox homeostasis. Physiol. Rev. 98, 1169–1203 (2018).

27. L. McInnes, J. Healy, J. Melville, UMAP: Uniform Manifold Approximation and Projection for Dimension Reduction, *arXiv [stat.ML]* (2018). http://arxiv.org/abs/1802.03426.

28. J. R. Giles, S. F. Ngiow, S. Manne, A. E. Baxter, O. Khan, P. Wang, R. Staupe, M. S. Abdel-Hakeem, H. Huang, D. Mathew, M. M. Painter, J. E. Wu, Y. J. Huang, R. R. Goel, P. K. Yan, G. C. Karakousis, X. Xu, T. C. Mitchell, A. C. Huang, E. J. Wherry, Shared and distinct biological circuits in effector, memory and exhausted CD8+ T cells revealed by temporal single-cell transcriptomics and epigenetics. Nat. Immunol. 23, 1600–1613 (2022).

29. D. Malhotra, E. Portales-Casamar, A. Singh, S. Srivastava, D. Arenillas, C. Happel, C. Shyr, N. Wakabayashi, T. W. Kensler, W. W. Wasserman, S. Biswal, Global mapping of binding sites for Nrf2 identifies novel targets in cell survival response through ChIP-Seq profiling and network analysis. Nucleic Acids Res. 38, 5718–5734 (2010).

30. A. Singh, S. Boldin-Adamsky, R. K. Thimmulappa, S. K. Rath, H. Ashush, J. Coulter, A. Blackford, S. N. Goodman, F. Bunz, W. H. Watson, E. Gabrielson, E. Feinstein, S. Biswal, RNAi-mediated silencing of nuclear factor erythroid-2-related factor 2 gene expression in non-small cell lung cancer inhibits tumor growth and increases efficacy of chemotherapy. Cancer Res. 68, 7975–7984 (2008).

31. A. Kopacz, D. Kloska, H. J. Forman, A. Jozkowicz, A. Grochot-Przeczek, Beyond repression of Nrf2: An update on Keap1. Free Radic. Biol. Med. 157, 63–74 (2020).

32. M. Komatsu, H. Kurokawa, S. Waguri, K. Taguchi, A. Kobayashi, Y. Ichimura, Y.-S. Sou, I. Ueno, A. Sakamoto, K. I. Tong, M. Kim, Y. Nishito, S.-I. Iemura, T. Natsume, T. Ueno, E. Kominami, H. Motohashi, K. Tanaka, M. Yamamoto, The selective autophagy substrate p62 activates the stress responsive transcription factor Nrf2 through inactivation of Keap1. Nat. Cell Biol. 12, 213–223 (2010).

33. Y. Inami, S. Waguri, A. Sakamoto, T. Kouno, K. Nakada, O. Hino, S. Watanabe, J. Ando, M. Iwadate, M. Yamamoto, M.-S. Lee, K. Tanaka, M. Komatsu, Persistent activation of Nrf2 through p62 in hepatocellular carcinoma cells. J. Cell Biol. 193, 275–284 (2011).

34. A. Kopacz, D. Klóska, B. Proniewski, D. Cysewski, N. Personnic, A. Piechota-Polańczyk, P. Kaczara, A. Zakrzewska, H. J. Forman, J. Dulak, A. Józkowicz, A. Grochot-Przęczek, Keap1 controls protein S-nitrosation and apoptosis-senescence switch in endothelial cells. Redox Biol. 28, 101304 (2020).

35. A.-M. Zavitsanou, R. Pillai, Y. Hao, W. L. Wu, E. Bartnicki, T. Karakousi, S. Rajalingam, A. Herrera, A. Karatza, A. Rashidfarrokhi, S. Solis, M. Ciampricotti, A. H. Yeaton, E. Ivanova, C. A. Wohlhieter, T. B. Buus, M. Hayashi, B. Karadal-Ferrena, H. I. Pass, J. T. Poirier, C. M. Rudin, K.-K. Wong, A. L. Moreira, K. M. Khanna, A. Tsirigos, T. Papagiannakopoulos, S. B. Koralov, KEAP1 mutation in lung adenocarcinoma promotes immune evasion and immunotherapy resistance. Cell Rep. 42, 113295 (2023).

36. A. Meister, Glutathione metabolism and its selective modification. J. Biol. Chem. 263, 17205–17208 (1988).

37. O. W. Griffith, Biologic and pharmacologic regulation of mammalian glutathione synthesis. Free Radic. Biol. Med. 27, 922–935 (1999).

38. A. C. Wild, H. R. Moinova, R. T. Mulcahy, Regulation of gamma-glutamylcysteine synthetase subunit gene expression by the transcription factor Nrf2. J. Biol. Chem. 274, 33627–33636 (1999).

39. T. Nguyen, P. Nioi, C. B. Pickett, The Nrf2-antioxidant response element signaling pathway and its activation by oxidative stress. J. Biol. Chem. 284, 13291–13295 (2009).

40. C. Tonelli, I. I. C. Chio, D. A. Tuveson, Transcriptional Regulation by Nrf2. Antioxid. Redox Signal. 29, 1727–1745 (2018).

41. G. Asantewaa, I. S. Harris, Glutathione and its precursors in cancer. Curr. Opin. Biotechnol. 68, 292–299 (2021).

42. E. Eruslanov, S. Kusmartsev, Identification of ROS using oxidized DCFDA and flow-cytometry. Methods Mol. Biol. 594, 57–72 (2010).

43. J.-C. Beltra, S. Manne, M. S. Abdel-Hakeem, M. Kurachi, J. R. Giles, Z. Chen, V. Casella, S. F. Ngiow, O. Khan, Y. J. Huang, P. Yan, K. Nzingha, W. Xu, R. K. Amaravadi, X. Xu, G. C. Karakousis, T. C. Mitchell, L. M. Schuchter, A. C. Huang, E. J. Wherry, Developmental relationships of four exhausted CD8+ T cell subsets reveals underlying transcriptional and epigenetic landscape control mechanisms. Immunity 52, 825–841.e8 (2020).

44. S. Moncada, J. R. Vane, Prostacyclin: its biosynthesis, actions and clinical potential. Philos. Trans. R. Soc. Lond. B Biol. Sci. 294, 305–329 (1981).

45. J. Stitham, C. Midgett, K. A. Martin, J. Hwa, Prostacyclin: An Inflammatory Paradox. Front. Pharmacol. 2 (2011).

46. D. Hanahan, Hallmarks of Cancer: New Dimensions. Cancer Discov. 12, 31–46 (2022).

47. J. Liu, Z. Chen, Y. Li, W. Zhao, J. Wu, Z. Zhang, PD-1/PD-L1 checkpoint inhibitors in tumor immunotherapy. Front. Pharmacol. 12, 731798 (2021).

48. H. O. Alsaab, S. Sau, R. Alzhrani, K. Tatiparti, K. Bhise, S. K. Kashaw, A. K. Iyer, PD-1 and PD-L1 checkpoint signaling inhibition for cancer immunotherapy: Mechanism, combinations, and clinical outcome. Front. Pharmacol. 8, 561 (2017).

49. J. Gong, A. Chehrazi-Raffle, S. Reddi, R. Salgia, Development of PD-1 and PD-L1 inhibitors as a form of cancer immunotherapy: a comprehensive review of registration trials and future considerations. J. Immunother. Cancer 6, 8 (2018).

50. C. J. Harvey, R. K. Thimmulappa, A. Singh, D. J. Blake, G. Ling, N. Wakabayashi, J. Fujii, A. Myers, S. Biswal, Nrf2-regulated glutathione recycling independent of biosynthesis is critical for cell survival during oxidative stress. Free Radic. Biol. Med. 46, 443–453 (2009).

51. V. B. Gubbala, N. Jytosana, V. Q. Trinh, H. C. Maurer, R. F. Naeem, N. K. Lytle, Z. Ma, S. Zhao, W. Lin, H. Han, Y. Shi, T. Hunter, P. K. Singh, K. P. Olive, M. C. B. Tan, S. M. Kaech, G. M. Wahl, K. E. DelGiorno, Eicosanoids in the pancreatic tumor microenvironment - a multicellular, multifaceted progression. Gastro Hep Adv. 1, 682–697 (2022).

52. A. E. Norlander, M. H. Bloodworth, S. Toki, J. Zhang, W. Zhou, K. Boyd, V. V. Polosukhin, J.-Y. Cephus, Z. J. Ceneviva, V. D. Gandhi, N. U. Chowdhury, L.-M. Charbonnier, L. M. Rogers, J. Wang, D. M. Aronoff, L. Bastarache, D. C. Newcomb, T. A. Chatila, R. S. Peebles Jr, Prostaglandin I2 signaling licenses Treg suppressive function and prevents pathogenic reprogramming. J. Clin. Invest. 131 (2021).

53. J. H. Chen, C. J. Perry, Y.-C. Tsui, M. M. Staron, I. A. Parish, C. X. Dominguez, D. W. Rosenberg, S. M. Kaech, Prostaglandin E2 and programmed cell death 1 signaling coordinately impair CTL function and survival during chronic viral infection. Nat. Med. 21, 327–334 (2015).

54. S. B. Lacher, J. Dörr, G. P. de Almeida, J. Hönninger, F. Bayerl, A. Hirschberger, A.-M. Pedde, P. Meiser, L. Ramsauer, T. J. Rudolph, N. Spranger, M. Morotti, A. J. Grimm, S. Jarosch, A. Oner, L. Gregor, S. Lesch, S. Michaelides, L. Fertig, D. Briukhovetska, L. Majed, S. Stock, D. H. Busch, V. R. Buchholz, P. A. Knolle, D. Zehn, D. Dangaj Laniti, S. Kobold, J. P. Böttcher, PGE2 limits effector expansion of tumour-infiltrating stem-like CD8+ T cells. Nature 629, 417–425 (2024).

55. M. Morotti, A. J. Grimm, H. C. Hope, M. Arnaud, M. Desbuisson, N. Rayroux, D. Barras, M. Masid, B. Murgues, B. S. Chap, M. Ongaro, I. A. Rota, C. Ronet, A. Minasyan, J. Chiffelle, S. B. Lacher, S. Bobisse, C. Murgues, E. Ghisoni, K. Ouchen, R. Bou Mjahed, F. Benedetti, N. Abdellaoui, R. Turrini, P. O. Gannon, K. Zaman, P. Mathevet, L. Lelievre, I. Crespo, M. Conrad, G. Verdeil, L. E. Kandalaft, J. Dagher, J. Corria-Osorio, M.-A. Doucey, P.-C. Ho, A. Harari, N. Vannini, J. P. Böttcher, D. Dangaj Laniti, G. Coukos, PGE2 inhibits TIL expansion by disrupting IL-2 signalling and mitochondrial function. Nature 629, 426– 434 (2024).

56. M. Cornberg, L. L. Kenney, A. T. Chen, S. N. Waggoner, S.-K. Kim, H. P. Dienes, R. M. Welsh, L. K. Selin, Clonal exhaustion as a mechanism to protect against severe immunopathology and death from an overwhelming CD8 T cell response. Front. Immunol. 4, 475 (2013).

57. I. Uboldi, P. Poduval, J. Prakash, “Engineering solutions to design CAR-T cells” in Engineering Technologies and Clinical Translation (Elsevier, 2022), pp. 1–31.

58. D. G. Roy, J. Chen, V. Mamane, E. H. Ma, B. M. Muhire, R. D. Sheldon, T. Shorstova, R. Koning, R. M. Johnson, E. Esaulova, K. S. Williams, S. Hayes, M. Steadman, B. Samborska, A. Swain, A. Daigneault, V. Chubukov, T. P. Roddy, W. Foulkes, J. A. Pospisilik, M.-C. Bourgeois-Daigneault, M. N. Artyomov, M. Witcher, C. M. Krawczyk, C. Larochelle, R. G. Jones, Methionine Metabolism Shapes T Helper Cell Responses through Regulation of Epigenetic Reprogramming. Cell Metab. 31, 250–266.e9 (2020).

59. W. Luo, M. S. Friedman, K. Shedden, K. D. Hankenson, P. J. Woolf, GAGE: generally applicable gene set enrichment for pathway analysis. BMC Bioinformatics 10, 161 (2009).

60. L. E. Dow, P. K. Premsrirut, J. Zuber, C. Fellmann, K. McJunkin, C. Miething, Y. Park, R. A. Dickins, G. J. Hannon, S. W. Lowe, A pipeline for the generation of shRNA transgenic mice. Nat. Protoc. 7, 374–393 (2012).

61. S. Nüssing, I. G. House, C. J. Kearney, A. X. Y. Chen, S. J. Vervoort, P. A. Beavis, J. Oliaro, R. W. Johnstone, J. A. Trapani, I. A. Parish, Efficient CRISPR/Cas9 gene editing in uncultured naive mouse T cells for in vivo studies. J. Immunol. 204, 2308–2315 (2020).

62. P. Heinrich, C. Kohler, L. Ellmann, P. Kuerner, R. Spang, P. J. Oefner, K. Dettmer, Correcting for natural isotope abundance and tracer impurity in MS-, MS/MS- and high-resolution-multiple-tracer-data from stable isotope labeling experiments with IsoCorrectoR. Sci. Rep. 8, 17910 (2018).

63. M. Martin, Cutadapt removes adapter sequences from high-throughput sequencing reads. EMBnet J. 17, 10 (2011).

64. A. Dobin, C. A. Davis, F. Schlesinger, J. Drenkow, C. Zaleski, S. Jha, P. Batut, M. Chaisson, T. R. Gingeras, STAR: ultrafast universal RNA-seq aligner. Bioinformatics 29, 15–21 (2013).

65. M. D. Robinson, D. J. McCarthy, G. K. Smyth, edgeR: a Bioconductor package for differential expression analysis of digital gene expression data. Bioinformatics 26, 139–140 (2010).

66. D. J. McCarthy, Y. Chen, G. K. Smyth, Differential expression analysis of multifactor RNA-Seq experiments with respect to biological variation. Nucleic Acids Res. 40, 4288– 4297 (2012).

67. Y. Chen, A. T. L. Lun, G. K. Smyth, From reads to genes to pathways: differential expression analysis of RNA-Seq experiments using Rsubread and the edgeR quasi-likelihood pipeline. F1000Res. 5, 1438 (2016).

68. T. Wu, E. Hu, S. Xu, M. Chen, P. Guo, Z. Dai, T. Feng, L. Zhou, W. Tang, L. Zhan, X. Fu, S. Liu, X. Bo, G. Yu, clusterProfiler 4.0: A universal enrichment tool for interpreting omics data. Innovation (Camb.) 2, 100141 (2021).

69. C. Schmidl, A. F. Rendeiro, N. C. Sheffield, C. Bock, ChIPmentation: fast, robust, low-input ChIP-seq for histones and transcription factors. Nat. Methods 12, 963–965 (2015).

