## Supplemental Figures for "NRF2-dependent regulation of the prostacyclin receptor PTGIR drives CD8 T cell exhaustion"

Dahabieh *et al.*

**This PDF file includes:**

Figs. S1-S7
Tables S1-S4

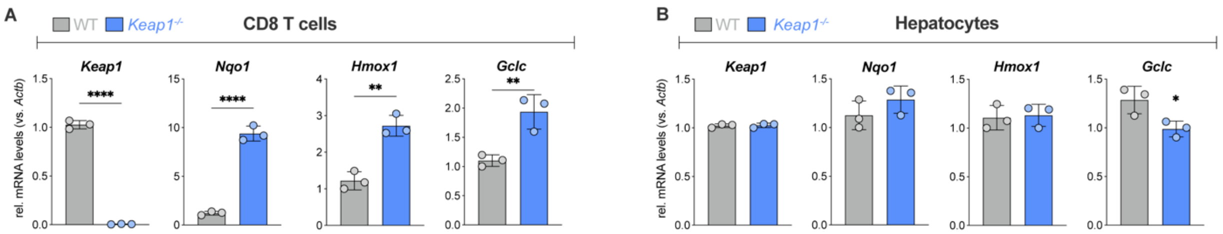


**Fig. S1. Validation of *Keap1* deletion in CD8 T cells. (A-B)** Relative mRNA expression for *Keap1*, and NRF2 transcriptional targets *Nqo1*, *Hmox1*, and *Gclc* in (A) CD8 T cells and (B) hepatocytes from $Keap1^{fl/fl}$ (WT) and $Cd4^{\text{Cre+}}$ $Keap1^{fl/fl}$ ($Keap1^{-/-}$) mice (mean±SEM, n=3). *$P$<0.05, **$P$<0.01, ****$P$<0.0001.

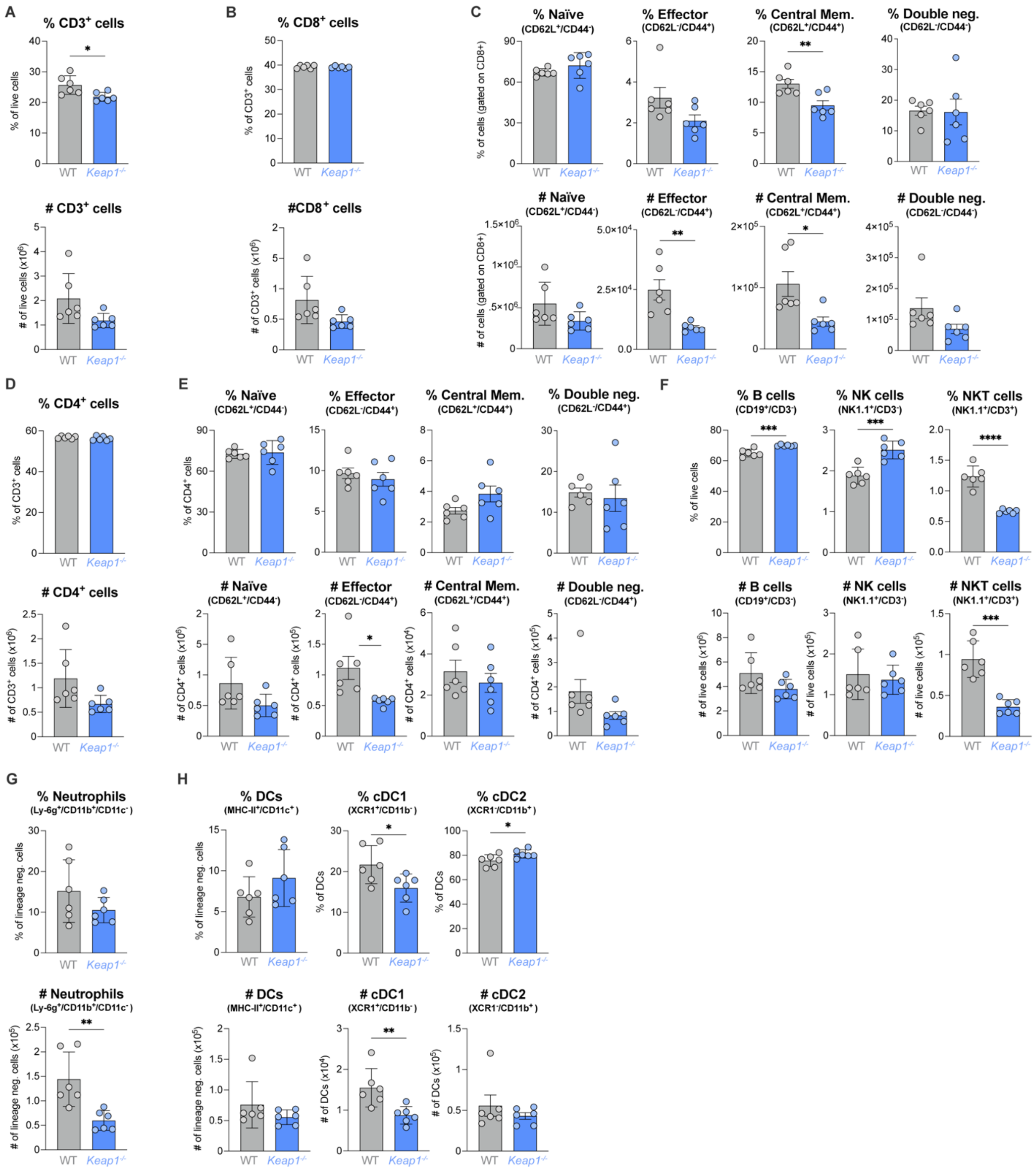


**Fig. S2. Immunophenotyping of splenocyte populations from *Keap1$^{fl/fl}$* (WT) and *Cd4*$^{Cre+}$ *Keap1$^{fl/fl}$* (*Keap1$^{-/-}$*) mice. (A-B)** Percent (top) and number (bottom) of splenic (A) CD3$^+$ T cells and **(B)** CD8$^+$ (gated on CD3$^+$) T cells from WT and *Keap1$^{-/-}$* mice aged 8-12 weeks (mean±SEM, n=6). **(C)** Percent (top) and number (bottom) of naïve, effector, central memory, and CD44$^-$

CD62L$^-$ (double negative) subsets of CD8$^+$ T cells in the spleens of mice as in (A). **(D-E)** CD4$^+$ T cells populations in WT and *Keap1*$^{-/-}$ mice. (D) Percent (top) and number (bottom) of CD4$^+$ (gated on CD3$^+$) T cells from WT and *Keap1*$^{-/-}$ mice aged 8-12 weeks (mean±SEM, n=6). **(E)** Percent (top) and number (bottom) of naïve, effector, central memory, and CD44$^-$CD62L$^-$ (double negative) subsets of CD4$^+$ T cells in the spleens of mice as in (D). **(F-H)** Percent (top) and number (bottom) of **(F)** B cells, Natural Killer (NK), and Natural Killer (NKT) T cells, **(G)** neutrophils, and **(H)** total CD11c$^+$ dendritic cells (DCs) and conventional DC1 (cDC1) and DC2 (cDC2) subsets from WT and *Keap1*$^{-/-}$ mice aged 8-12 weeks (mean±SEM, n=6). *$P$<0.05, **$P$<0.01, ***$P$<0.001, ****$P$<0.0001.

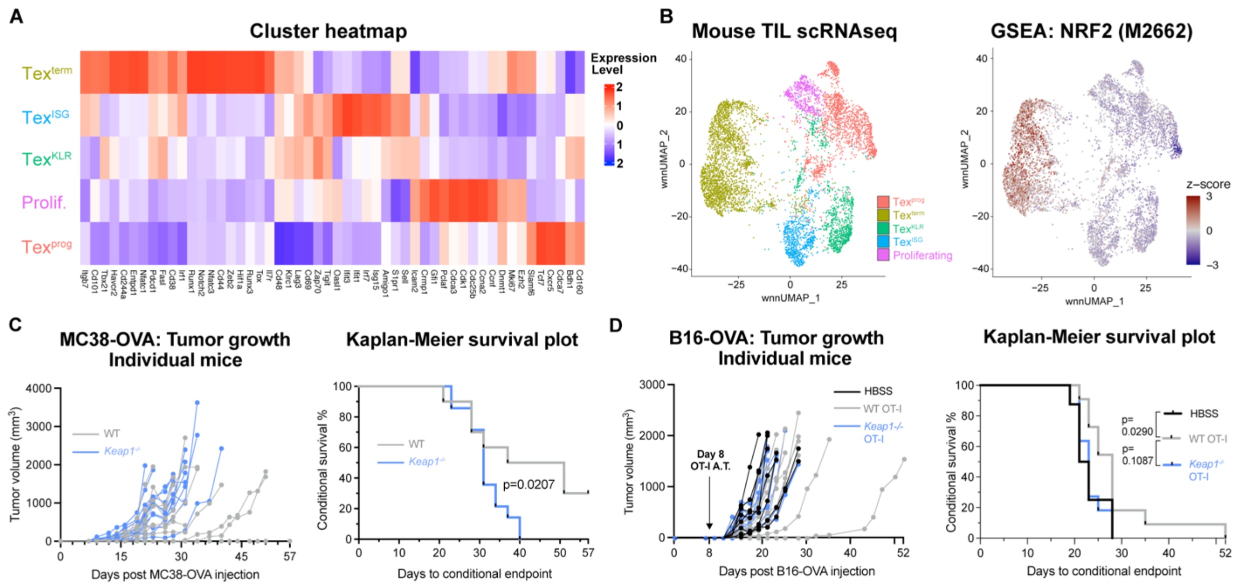


**Fig. S3. scRNA-seq cluster analysis and impact of *Keap1* deletion in T cells on tumor growth and survival. (A)** Heatmap of gene expression markers for progenitor exhausted (Tex$^{prog}$), terminally exhausted (Tex$^{term}$), exhausted killer cell lectin-like receptor-expressing (Tex$^{KLR}$), exhausted IFN-I stimulated gene expressing (Tex$^{ISG}$)**,** and proliferating (prolif) CD8$^{+}$ TIL isolated from B16-OVA tumors (related to Figure 2A). **(B)** Weighted nearest neighbor UMAP for CD8 TIL clusters described in (A) and embedding of GSEA for NRF2 transcriptional target signature (M2662) across clusters. **(C)** Individual growth curves for MC38-OVA tumors in WT and *Keap1*$^{-/-}$ mice. *Right*, Kaplan-Meier (KM) survival curves for time to tumor endpoint (>1500 mm$^3$) for MC38-OVA tumors grown in WT and *Keap1*$^{-/-}$ mice. **(D)** Individual growth curves for B16-OVA tumors following adoptive transfer of WT or *Keap1*$^{-/-}$ OT-I CD8 T cells at 8 dpti. *Right*, Kaplan-Meier (KM) survival curves for time to tumor endpoint (>1500 mm$^3$) for B16-OVA tumors.

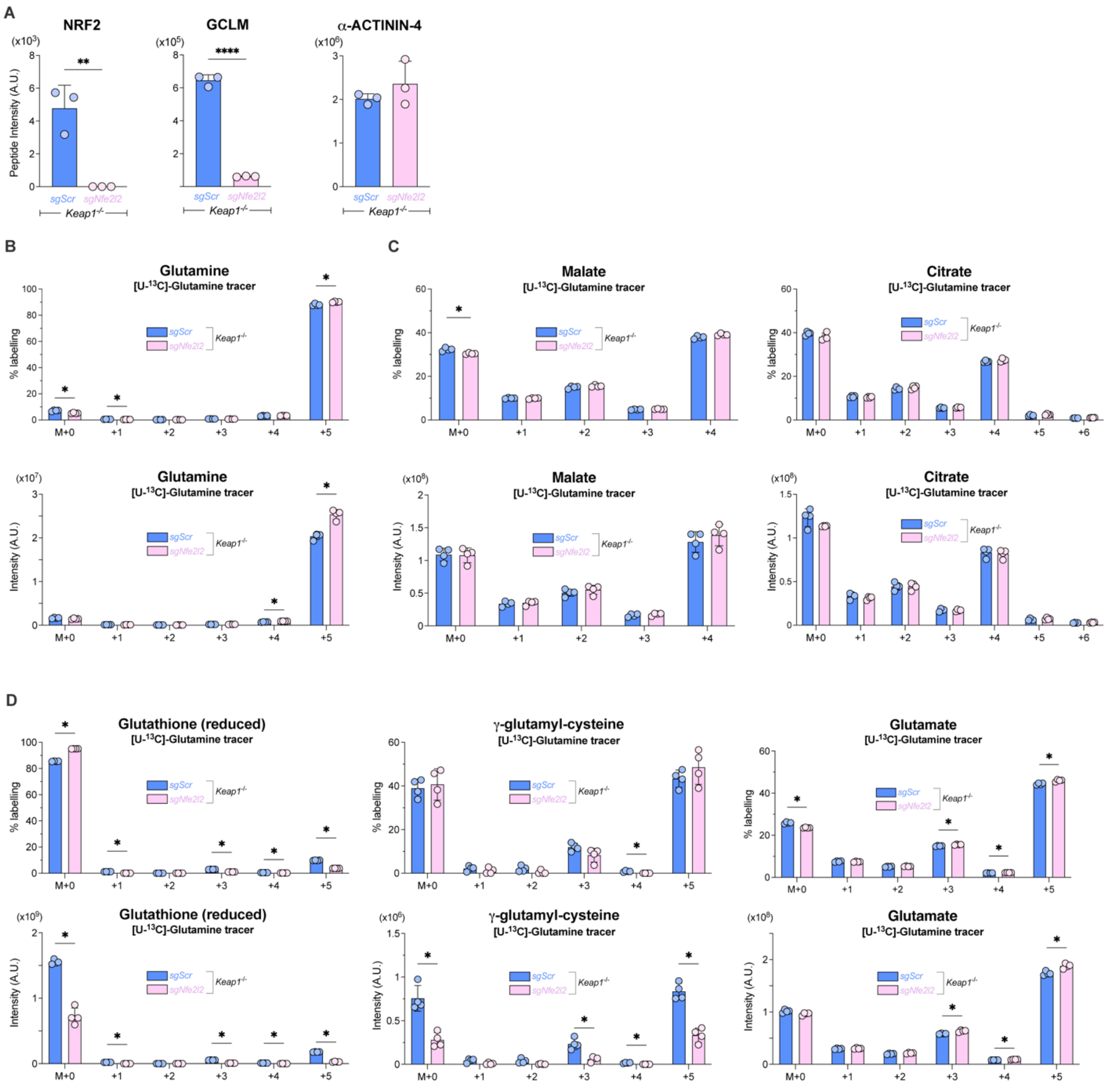


**Fig. S4. Proteomic and metabolic profiling of *Keap1*$^{-/-}$ CD8 T cells. (A)** Raw peptide intensities for NRF2, GCLM, and ACTN4 in *Keap1*$^{-/-}$ CD8$^{+}$ P14 T cells that received *Nfe2l2*-targeting (*sgNfe2l2*) or scrambled control (*sgScr*) sgRNAs (mean±SEM, n=3). **(B-D)** Mass isotopologue distribution (MID) for intracellular metabolites from *sgNfe2l2* and *sgScr Keap1*$^{-/-}$ P14 T cells following culture with [U-$^{13}$C]-Glutamine (mean±SEM, n=4). Shown are the percent labelling (top) and intensities (bottom) for indicated isotopologues for **(B)** Glutamine, **(C)** Malate and

Citrate, and **(D)** Glutathione (reduced), γ-glutamyl cysteine, and Glutamate. $*P<0.05$, $**P<0.01$, $****P<0.0001$.

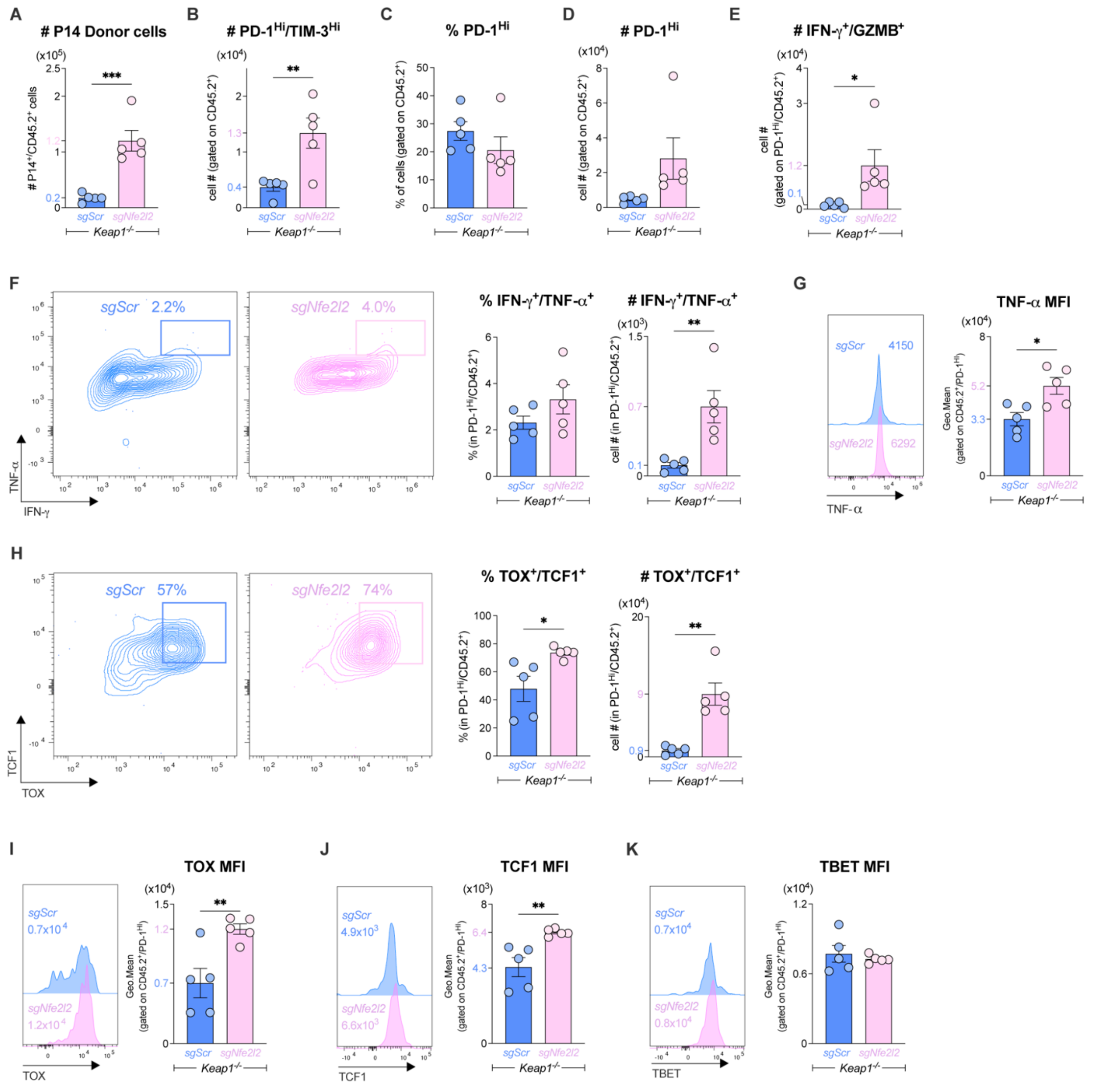


**Fig. S5. Immunophenotyping of *Keap1$^{-/-}$ sgScr* and *sgNfe2l2* P14 cells following LCMV CL13 infection. (A-E)** Bar graphs quantifying **(A)** the number of P14 (CD45.2$^{+}$) donor cells, **(B)** number of PD-1$^{Hi}$TIM-3$^{Hi}$ donor P14 cells, **(C)** percentage of PD-1$^{Hi}$ donor P14 cells, **(D)** number of PD-1$^{Hi}$ donor P14 cells, and **(E)** number of IFN-γ$^{+}$GZMB$^{+}$ P14 cells from the spleen of LCMV CL13-infected mice (7 dpi) that received *sgNfe2l2* or *sgScr Keap1$^{-/-}$* P14 (CD45.2$^{+}$) donor T cells at -1 dpi (mean±SEM, n=5). **(F)** Quantification of the percentage and number of polyfunctional (IFN-

$\gamma^+$TNF-$\alpha^+$) PD-$1^{Hi}$ *sgNfe2l2* or *sgScr Keap1*$^{-/-}$ P14 CD8 T cells from the spleen of LCMV CL13-infected mice (7 dpi). **(G)** Representative histograms and bar graph of MFI for TNF-$\alpha$ expression by PD-$1^{Hi}$ donor cells from (F) at 7 dpi (mean±SEM, n=5/sample). **(H)** Quantification of the percentage and number of TOX$^+$TCF1$^+$ PD-$1^{Hi}$ *sgNfe2l2* or *sgScr Keap1*$^{-/-}$ P14 CD8 T cells from the spleen of LCMV CL13-infected mice at 7 dpi (mean±SEM, n=5). **(I-K)** Representative histograms and bar graph of MFI for (I) TOX, (J) TCF1, and (K) TBET expression by PD-$1^{Hi}$ donor cells from (F) at 7 dpi (mean±SEM, n=5/sample).*$P<0.05$, **$P<0.01$, ***$P<0.001$.

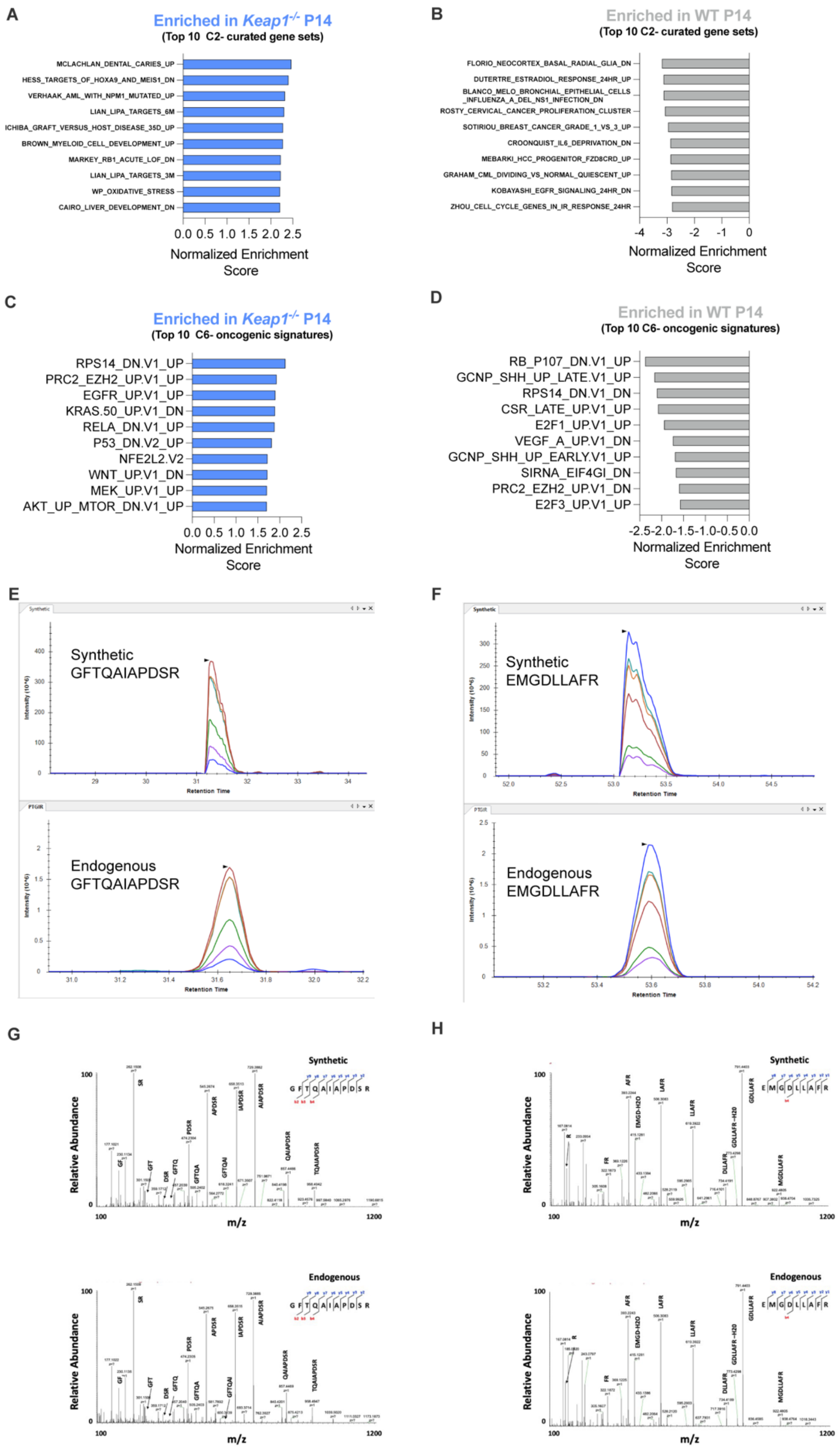

A
Enriched in Keap1-/- P14
(Top 10 C2- curated gene sets)
MCLACHLAN_DENTAL_CARIES_UP
HESS_TARGETS_OF_HOXA9_AND_MEIS1_DN
VERHAAK_AML_WITH_NPM1_MUTATED_UP
LIAN_LIPA_TARGETS_6M
ICHIBA_GRAFT_VERSUS_HOST_DISEASE_35D_UP
BROWN_MYELOID_CELL_DEVELOPMENT_UP
MARKEY_RB1_ACUTE_LOF_DN
LIAN_LIPA_TARGETS_3M
WP_OXIDATIVE_STRESS
CAIRO_LIVER_DEVELOPMENT_DN
0.0 0.5 1.0 1.5 2.0 2.5
Normalized Enrichment Score
B
Enriched in WT P14
(Top 10 C2- curated gene sets)
FLORIO_NEOCORTEX_BASAL_RADIAL_GLIA_DN
DUTERTRE_ESTRADIOL_RESPONSE_24HR_UP
BLANCO_MELO_BRONCHIAL_EPITHELIAL_CELLS_INFLUENZA_A_DEL_NS1_INFECTION_DN
ROSTY_CERVICAL_CANCER_PROLIFERATION_CLUSTER
SOTIRIOU_BREAST_CANCER_GRADE_1_VS_3_UP
CROONQUIST_IL6_DEPRIVATION_DN
MEBARKI_HCC_PROGENITOR_FZD8CRD_UP
GRAHAM_CML_DIVIDING_VS_NORMAL_QUIESCENT_UP
KOBAYASHI_EGFR_SIGNALING_24HR_DN
ZHOU_CELL_CYCLE_GENES_IN_IR_RESPONSE_24HR
-4 -3 -2 -1 0
Normalized Enrichment Score
C
Enriched in Keap1-/- P14
(Top 10 C6- oncogenic signatures)
RPS14_DN.V1_UP
PRC2_EZH2_UP.V1_UP
EGFR_UP.V1_UP
KRAS.50_UP.V1_DN
RELA_DN.V1_UP
P53_DN.V2_UP
NFE2L2.V2
WNT_UP.V1_DN
MEK_UP.V1_UP
AKT_UP_MTOR_DN.V1_UP
0.0 0.5 1.0 1.5 2.0 2.5
Normalized Enrichment Score
D
Enriched in WT P14
(Top 10 C6- oncogenic signatures)
RB_P107_DN.V1_UP
GCNP_SHH_UP_LATE.V1_UP
RPS14_DN.V1_DN
CSR_LATE_UP.V1_UP
E2F1_UP.V1_UP
VEGF_A_UP.V1_DN
GCNP_SHH_UP_EARLY.V1_UP
SIRNA_EIF4GI_DN
PRC2_EZH2_UP.V1_DN
E2F3_UP.V1_UP
-2.5-2.0-1.5-1.0-0.5 0.0
Normalized Enrichment Score
E
Synthetic GFTQAIAPDSR
Intensity (10^6)
Retention Time
Endogenous GFTQAIAPDSR
F
Synthetic EMGDLLAFR
Endogenous EMGDLLAFR
G
Synthetic
Endogenous
Relative Abundance
m/z
100
1200
H
Synthetic
Endogenous
Relative Abundance
m/z

**Fig. S6. Enriched gene sets in *Keap1$^{-/-}$* versus WT P14 cells and mass spectrometry validation of PTGIR peptides. (A-D)** Top 10 enriched pathways for (**A-B**) MSigDB C2 (curated) gene sets and (**C-D**) MSigDB C6 (oncogenic signature) gene sets in *Keap1$^{-/-}$* versus WT P14 T cells following LCMV infection (7 dpi). Data are plotted as Normalized Enrichment Score (NES) for each pairwise comparison. **(E-F)** Representative PRM transition traces for the synthetic (top) and endogenous (bottom) PTGIR peptides (**E**) GFTQAIAPDSR and **(F)** EMGDLLAFR. **(G-H)** Representative MS2 spectra of the synthetic (top) and endogenous (bottom) peptides (**G**) GFTQAIAPDSR and **(H)** EMGDLLAFR of PTGIR.

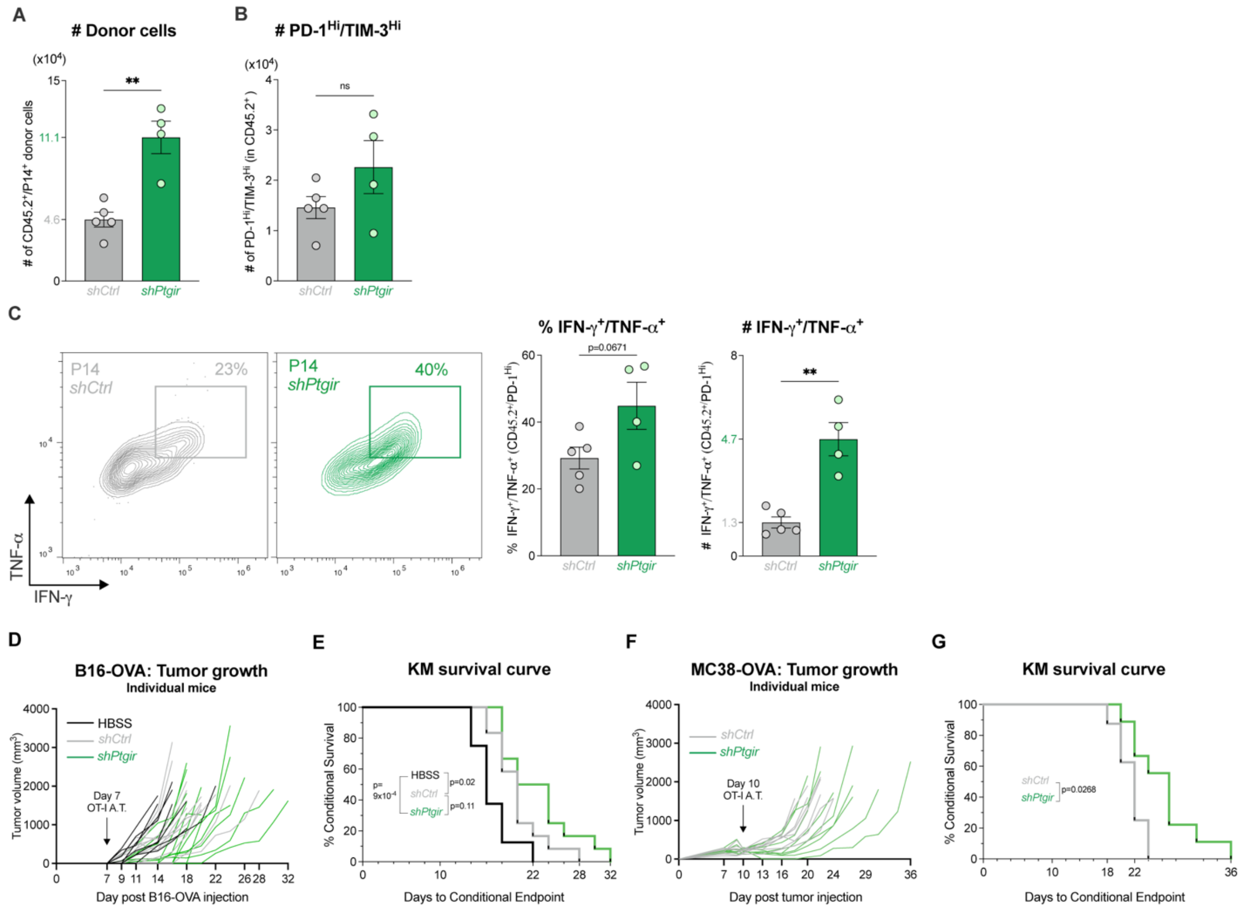


**Fig. S7. *Ptgir* knockdown improves anti-viral CD8 T cell responses and attenuates tumor growth. (A)** Bar graph quantifying the number of control (*shCtrl*) and *Ptgir* knockdown *(shPtgir)* P14 donor cells in the spleen of LCMV CL13-infected mice at 8 dpi (mean±SEM, n=4-5/sample). CD45.2 is a donor cell marker expressed by P14 T cells. **(B)** Bar graph of PD-$1^{Hi}$TIM-$3^{Hi}$ *shCtrl*-versus *shPtgir*-expressing P14 T cells from cells in (A). **(C)** Quantification of the percentage and number of polyfunctional (IFN-$\gamma^+$TNF-$\alpha^+$) PD-$1^{Hi}$ *shCtrl*- versus *shPtgir*-expressing P14 T donor cells from LCMV CL13-infected mice at 8 dpi (mean±SEM, n=4-5/sample). **(D)** Individual growth curves for B16-OVA tumors following adoptive transfer of OT-I CD8 T cells expressing control (*shCtrl*) and *Ptgir (shPtgir)* shRNAs at 7 dpti. *Right*, Kaplan-Meier (KM) survival curves for time to tumor endpoint (>1500 $mm^3$) for B16-OVA tumors. **(E)** Individual growth curves for MC38-

OVA tumors following adoptive transfer of OT-I CD8 T cells expressing control (*shCtrl*) and *Ptgir (shPtgir)* shRNAs at 10 dpti. *Right*, Kaplan-Meier (KM) survival curves for time to tumor endpoint (>1500 $mm^3$) for MC38-OVA tumors.**P*<0.05, ***P*<0.01.

**Table S1: Gene Set Enrichment Analysis (GSEA) of the oncogenic signature gene sets (MSigDB C6 gene set) enriched in Tex versus Teff cell clusters.** RNA-seq data are mined from GSE89307, GSE84820, and GSE86881. Refer to first tab of excel sheet for detailed description of column headings in subsequent data tab.

**Table S2: qPCR, shRNA and sgRNA oligonucleotide sequences.** 5' to 3' oligonucleotide sequences used for qPCR, CRISPR-Cas9 mediated knockout, and shRNA-mediated knockdown. Q denotes primers used for qPCR, and F and R denote forward and reverse primers, respectively.

**Table S3: Differential gene expression analysis from RNA-seq of adoptively transferred *Keap1$^{-/-}$* versus WT P14 cells, isolated 7 days post LCMV Armstrong infection.** Refer to first tab of excel sheet for detailed description of column headings in subsequent data tab.

**Table S4: C2, C5, C6, C7, NFE2L2.v2 and H Gene set enrichment analysis (GSEA) from RNA-seq of *Keap1$^{-/-}$* versus WT P14$^+$ CD8$^+$ T cells upon LCMV Armstrong infection.** A Positive or negative normalized enrichment score (NES) indicates a gene set enrichment in *Keap1$^{-/-}$* cells or WT cells, respectively. Refer to first tab of excel sheet for detailed description of column headings in subsequent data tabs.
